# Phase-separating enzyme-responsive condensates for inhalable delivery of biomacromolecules

**DOI:** 10.1101/2025.06.17.660252

**Authors:** Yang Xiao, Shijia Zhou, Ting Zhu, Yulu Du, Qiyong Hu, Xinyi Guo, Long Wang, Shengyuan Deng, Kewei Ren

## Abstract

Inhaled delivery of biomacromolecules shows extensive promise for treating a large variety of diseases, especially pulmonary and respiratory diseases. Although nanoparticle-based vehicles enhance intracellular uptake efficiency, it still remains challenging for biomacromolecules inhaled delivery to overcome cytotoxicity, cargo leakage and off-target effect. Here, we present an enzyme-responsive condensates based highly biocompatible and controllable platform for inhaled delivery of biomacromolecules. The condensates that can be modularly designed for rapid recruitment and enrichment of biomolecules, including Cas9 ribonucleoprotein (RNP) and mRNA, were internalized by cells via lysosomal-independent endocytosis, and subsequently released payloads triggered with endogenous physicochemical stimulus for controlled regulation of genes and proteins expression. Additionally, the micrometer sized condensates with highly structural and biological stability during lyophilization and nebulization enable inhalable delivery of biomacromolecules to lung. This condensate-based vehicle provides a promising platform for biomacromolecules inhaled delivery and controlled release, targeted cell regulation, and precise therapy.

## 1. Introduction

With high efficacy,[1] specificity[2] and biocompatibility,[3] biomacromolecules, such as proteins and RNAs[4] have emerged as powerful therapeutic agents for several diseases treatment. Efficient and safe delivery of exogenous biomacromolecules into target cells is crucial to their therapeutic application.[5–8] Currently, various vehicles including viral vectors,[9] lipid[10,11] and polymers nanoparticles,[12–14] have been used to deliver biomacromolecules into cells. Although these systems demonstrated the promising application for biomacromolecules transportation, they also have drawbacks, including potential cytotoxicity, low loading efficiency, fabrication complexity and poor endosomal escape.[15–18] Consequently, new vehicles capable of achieving efficient and biocompatible biomacromolecule delivery are still urgently needed.

Biomolecular condensates that arisen from liquid-liquid phase separation can recruit and condense, such as proteins and RNAs from a homogeneous state to a concentrated state.[19,20] Due to the benefits of negligible cytotoxicity, fast and efficient drugs recruitment ability, specifically localized environment to maintain payloads bioactivity, and endocytosis-independent cellular internalization pathway, biomolecular condensates have opened new avenues for delivering functional biomolecules within cells.[8,21] For examples, Gu and co-workers constructed a coacervate vesicle by using of DNA and histones for biopharma-ceuticals intracellular delivery.[22] Peptides based coacervates have been designed for cytosolic delivery and redox-activated release of DNA, RNA and proteins.[8,21] However, these condensates-based delivery vehicles were administered by traditional injection, the application of biomolecular condensates for inhaled drug delivery have yet to be developed. In this study, we will introduce a new type of inhalable condensate system to fill this gap.

Compared to injection-based administration, inhalation preparations offers several advantages such as noninvasiveness,[23] higher local drug accumulation,[24] lower systemic side effects,[25] and reduced nonspecific interaction between drugs and serum proteins.[26] Although several carriers such as lipid nanoparticle, inorganic nanoparticles and virus-based systems[27] are recently developed for inhaled delivery of biomacromolecules, especially mRNAs and proteins, the major limitations of cargo leakage, potential off-target effects and toxicity persist as concerns.[28,29] In addition, to achieve desired therapeutic efficacy, the stimuli-responsive and “on-demand” drug release is extremely desired in inhaled delivery system.[30]

Here, we present a novel peptide-based condensate systems for inhalable delivery and protein kinase A (PKA)-responsive release of biomacromolecules. We demonstrate that the condensates are capable of cellular uptake via lysosomal independent endocytic pathway for efficient delivery of various biomacromolecules of Cas9 ribonucleoprotein (RNP) or mRNA, which shows lower cytotoxicity and higher transfection efficiency than commercial transfection reagents. In addition, the condensates can be modularly designed, easily synthesized and achieved endogenous mRNA and PKA cooperatively controlled RNP activation and gene editing. We show that the micrometer-sized condensates exhibited highly structural and biological stability, and facilitated dry powder aerosol delivery to lung (**Figure 1**). Overall, these enzyme-responsive condensates can be used as general platform for a broad range of biomacromolecules inhalable delivery, controlled release, and precise gene therapy.

**Figure 1.**
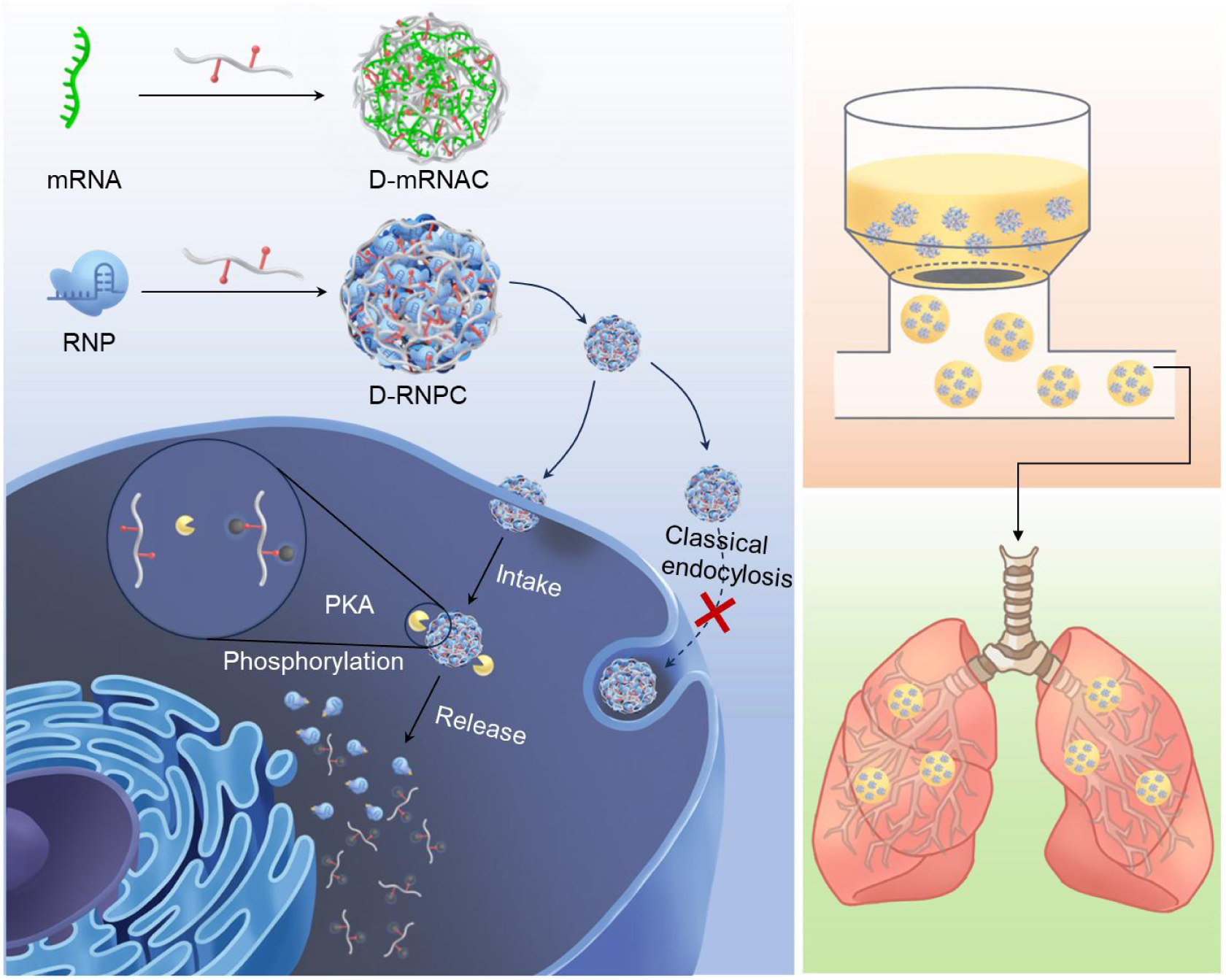
Schematic illustration of enzyme-responsive condensates design for cytoplasmic and inhalable delivery of biomacromolecules. Biomacromolecules of ribonucleoprotein (RNP) or messenger RNA (mRNA) are recruited and formed condensate by LLPS. The condensate can be uptaken by cells via a lysosomal-independent endocytic pathway, resulting in the intracellular PKA triggered disassembly and release of the therapeutic for regulating gene and protein expression. Condensate-mediated biomacromolecules delivery into lung after nebulization administration.

## 2. Results and Discussion

### 2.1. Synthesis and Characterization of Dual Peptide/sgRNA/Cas9 Condensate (D-RNPC)

We chose the dual peptide sequence RRASLRRASL, which can be phosphorylated by PKA,[31] as a polycation to form condensates of biomacromolecules such as proteins and mRNA through electrostatic interaction. The dual peptide was labelled with 5-carboxyfluorescein (FAM) dye, and Cas9 was labelled with red fluorescent protein (RFP). Confocal laser scanning microscopy (CLSM) imaging revealed that the single component of Cas9 or dual-peptide (D-Pep) was uniformly distributed in the solution, and the Cas9 and sgRNA complex (RNP) formed nanoscale particle. After the dual peptide was mixed with RNP, the spherical-shaped biomolecular condensates with micron size were observed (Figure 2a). The fluorescence signals of the FAM and RFP in dual peptide/sgRNA/Cas9 condensates (D-RNPC) were overlapped, and exhibited a position-sensitive dependence (Figure S1, Supporting Information). Z-stack images of D-RNPC demonstrated that the Cas9 and dual peptide were distributed inside the condensates (Figure S2, Supporting Information). However, the dual peptide and positively charged Cas9 mixture (D-PC) failed to form the condensates, indicating the LLPS was driven by electrostatic interactions. To evaluate whether the triple peptide, RRASLRRASLRRASL, that possesses a stronger positive charge, enable to enhance the loading capacity of RNP, we prepared the triple peptide/sgRNA/Cas9 condensates (T-RNPC) (Figure S3, Supporting Information).

**Figure 2.**
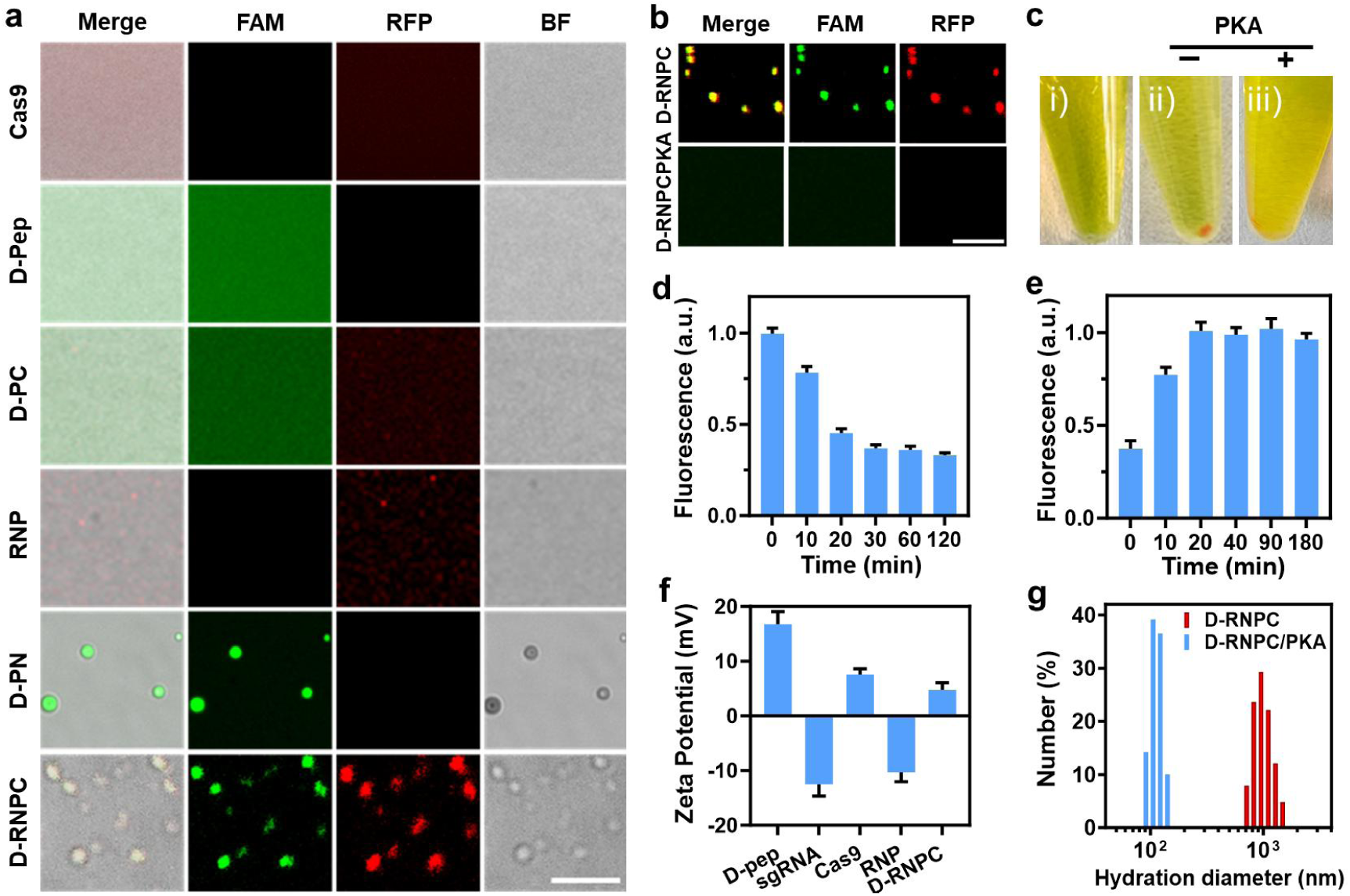
Characterization of D-RNPC preparation and decomposition. a) Confocal fluorescence imaging of D-RNPC, dual-peptide and sgRNA mixture (D-PN), RNP, the dual peptide and positively charged cas9 mixture (D-PC), dual-peptide (D-Pep) and Cas9. Scale bar, 5 µm. b) Confocal fluorescence imaging of 200 nM D-RNPC in absence (D-RNPC) and presence of 1400 nM PKA (D-RNPC/PKA). Scale bar, 5 µm. c) Morphology of i) mixing of dual peptide and RNP then immediate centrifugation, ii) 200 nM D-RNPC after centrifugation, iii) incubation of 1400 nM PKA and 200 nM D-RNPC for 1 h, then centrifugation. d) Fluorescence statistics of the supernatant during the formation of D-RNPC. Shown are mean ± SEM (n = 3). e) Fluorescence statistics of the supernatant during the PKA-triggered D-RNPC decomposition. Shown are mean ± SEM (n = 3). f) Zeta potentials of D-pep, sgRNA, Cas9, RNP and. D-RNPC. Shown are mean ± SEM (n = 3). g) Hydrodynamic diameter distribution of 200 nM D-RNPC, and mixture of 200 nM D-RNPC and 1400 nM PKA after 1-h incubation.

We first investigated the effect of peptide concentration on formation of condensates. With the increasing of peptides, the amounts of D-RNPC and T-RNPC were increased until the dual peptide and triple peptide concentration reach to 200 µM and 75 µM (Figure S4A, Supporting Information), respectively, which were selected to conduct the subsequent experiments. In addition, the optimal ratio of sgRNA to Cas9 for D-RNPC or T-RNPC preparation was 1:1 (Figure S4B, Supporting Information). After incubation in Minimal Essential Medium (MEM) and physiological saline (NS) for 1 h, the size and amount of D-RNPC and T-RNPC showed almost no change (Figure S5A, Supporting Information). Moreover, the negligible influence of temperature on condensate formation was observed (Figure S5B, Supporting Information). All above results demonstrated the high stability of D-RNPC and T-RNPC.

Upon addition of Protein Kinase A (PKA), the phosphorylation of dual peptide results in a reduction of positive charge density, leading to the dissolution of D-RNPC and the release of RNP (Figure 2b,c). The PKA-dependent disassembly of D-RNPC was achieved after addition of PKA with different concentrations (Figure S6, Supporting Information). Confocal microscopy imaging demonstrated that after 1 h of PKA treatment, about 90% of D-RNPC was disassembled. In contrast, only approximately 40% of T-RNPC was disassembled. In addition, the M-RNPC assembled with RNP and a misorder peptide, RSRALRSRAL, that cannot be phosphorylated by PKA, remained stable and almost unable to be decomposed upon the addition of PKA (Figure S7, Supporting Information), exhibiting the high specificity and selectivity of PKA.

To further test condensates formation and PKA activated disassembly, we mixed the peptides and RNP in the buffer solution (FIGUREREF 2c) and incubated for different times, then measured the fluorescence intensity of supernatant after centrifugation. After incubation of D-RNPC for 1 h and centrifugation, the precipitation with red fluorescent signal was observed at the bottom of tube, which disappeared in presence of PKA (Figure 2c). As the incubation time between peptides and RNP increased, the fluorescence intensity of FAM in supernatant gradually decreased (Figure 2d; Figure S8A, Supporting Information), indicating that the 5-FAM labeled peptides progressively aggregate into the condensate. Similarly, the fluorescence intensities in supernatant of M-RNPC and T-RNPC also reduced over time (Figure S8C, E, Supporting Information). Upon addition of PKA, the fluorescence intensity of the D-RNPC supernatant was enhanced (Figure 2e; Figure S8B, Supporting Information), suggesting the decomposition of D-RNPC and release of 5-FAM labeled peptides. Compared to the D-RNPC, the decomposition rate of T-RNPC induced by PKA was slower (Figure S8D, Supporting Information). Therefore, D-RNPC was used for the subsequent experiments. As a control, no obvious fluorescence intensity enhancement was observed in the supernatant of M-RNPC (Figure S8F, Supporting Information). All these results suggested the high specificity and efficiency of PKA for dual peptide phosphorylation.

Under the optimal conditions, the D-RNPC was prepared, which resulted in approximately 50-fold enrichment of Cas9 (Figure S9, Supporting Information). Zeta potential analysis also further verified the formation of D-RNPC (Figure 2f), and the size of D-RNPC was around 1 µm. The D-RNPC could be disassembled into 110 nm upon addition of PKA (Figure 2g), which is consistent with RNP condensates size (Figure S10, Supporting Information). D-RNPC remained stable even after incubation for 72 h (Figure S11, Supporting Information)

### 2.2. Cellular Uptake and PKA Activated Release of D-RNPC

To evaluate the genome editing ability of D-RNPC, Polo-like kinase 1 (*PLK1*) gene was selected as a model therapeutic target, which was extracted from human cervical cancer (HeLa) cells and served as a template for PCR. Gel electrophoresis analysis demonstrated that the RNP could effectively cleave the target DNA generated with PCR (Figure S12, Supporting Information).

We subsequently tested the application of D-RNPC in living cells. As shown in Figure S13, Supporting Information, the RFP signals of Cas9 overlapped well with DAPI, indicating the D-RNPC could be efficiently uptaken by HeLa cells, and intracellular over-expressed PKA induced the release of RNP for nuclear delivery. As control, the fluorescence of Cas9 was observed on the cell membrane but not inside of cells incubated with RNP, Cas9 or the Cas9 and dual peptide mixture (D-PC), demonstrating that only Cas9 formed into condensate could enable cellular internalization. Next, we investigated the influences of incubation time on cellular internalization and disassembly of D-RNPC. The intracellular fluorescence signal of FAM was in an incubation time-dependent manner and reached saturation at 12 h, while the lowest Pearson’s correlation value between the signals of FAM and RFP was achieved at 24 h, suggesting the optimal incubation times for cellular internalization and decomposition of D-RNPC (Figure S14, Supporting Information).

To demonstrate that PKA specifically triggered the dissociation of D-RNPC, the PKA inhibitor H-89 was used to decrease the PKA expression in HeLa cells. Compared with HeLa cells incubated with D-RNPC, the higher overlaps of FAM and RFP signals were observed in HeLa cells treated with M-RNPC, or H-89 and D-RNPC (Figure 3a; Figure S15A, Supporting Information). While the maximum overlap of DAPI and RFP signals with Pearson’s correlation value of 0.4 was observed in cells incubated with D-RNPC (Figure 3a; Figure S15B, Supporting Information). All these results demonstrated the high specificity of PKA activated release and nucleus delivery of RNP. Similarly, the negligible release and nuclear delivery of RNP were exhibited in α-mouse liver 12 (AML-12) cells with lower PKA content (Figure 3b; Figure S16, Supporting Information), demonstrating the cell selective release of D-RNPC.

**Figure 3.**
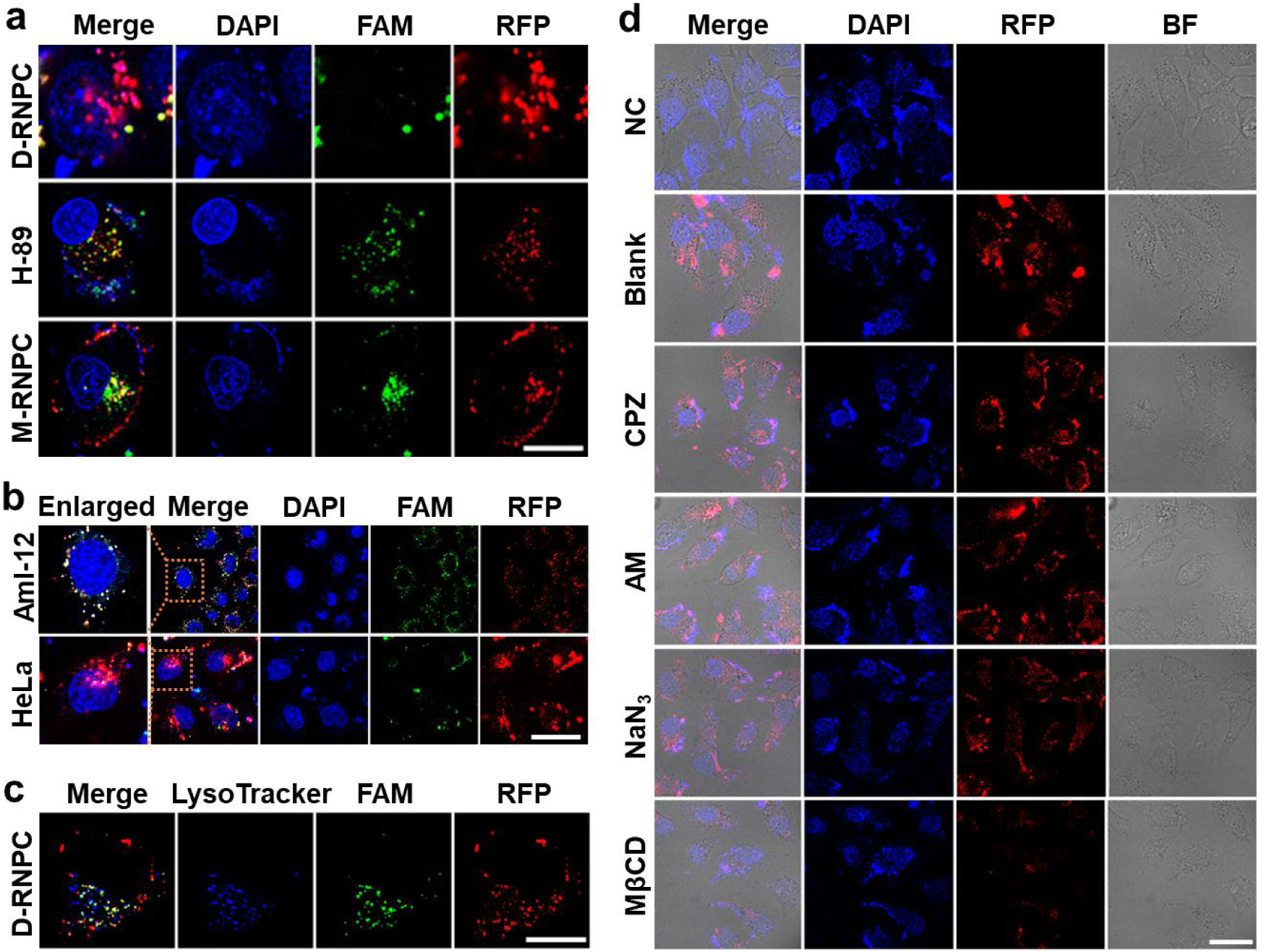
Cellular internalization of D-RNPC. a) Confocal fluorescence imaging of HeLa cells incubated with 200 nM D-RNPC in absence (D-RNPC) and presence of 20 μM H-89 (H-89), and 200 nM M-RNPC for 24 h. Scale bar, 25 µm. b) Confocal fluorescence imaging of HeLa cells and AML-12 cells incubated with 200 nM D-RNPC for 24 h. Scale bar, 100 µm. c) Confocal fluorescence imaging of HeLa cells incubated with 200 nM D-RNPC for 12 h. Scale bar, 25 µm. d) Confocal fluorescence imaging uptake inhibition of 200 nM D-RNPC in HeLa cells preincubated without treatment (Blank) and with methyl-β-cyclodextrin (MeβCD, 2.5 mM), sodium azide (NaN₃, 100 mM), amiloride (AM, 20 μM) and chlorpromazine (CPZ, 30 μM). The cells without treatment as negative control (NC). Scale bar, 25 µm.

To investigate the lysosome escape capability of D-RNPC, LysoTracker Blue was used to stain acidic organelles. As shown in Figure 3c, a poor colocalization between lysosomes and D-RNPC was observed in HeLa cells, verifying the successful lysosomal escape of D-RNPC. The internalization pathway of the D-RNPC in HeLa cells was further studied by using various inhibitors. The cellular uptake of D-RNPC was inhibited approximately 50% by Methyl-β-cyclodextrin (MβCD, a cholesterol-depleting reagent). In comparison, the negligible fluorescence signal decrease was observed in cells pretreated by chlorpromazine (CPZ, an inhibitor of clathrin-dependent pathways), amiloride (AM, an inhibitor of macropinocytosis), and sodium azide (NaN₃, an inhibitor of energy-dependent endocytosis) (Figure 3d; Figure S17, Supporting Information), which is consistent with the results from a flow cytometry assay (Figure S18, Supporting Information). All these results suggested that the mechanism of D-RNPC uptake is lipid raft-mediated endocytosis, and not lysosomal-dependent endocytosis.[32,33] In addition, D-RNPC demonstrated higher capabilities (Figure S19, Supporting Information) and biocompatibility (Figure S20, Supporting Information) for RNP delivery than the commercial transfection reagents.

### 2.3. D-RNPC Mediated Target Gene Editing and Expression Inhibition in Living Cells

To further assess the gene cleavage efficiency of D-RNPC in HeLa cells, T7 endonuclease 1 (T7E1) assay was performed. The results of agarose gel electrophoresis exhibited that about 42.6% target genes were knocked down in HeLa cells treated with D-RNPC, whereas almost no gene cleavage was observed in cells incubated with M-RNPC (Figure 4a). Subsequently, we evaluated the inhibitory efficiency of D-RNPC on intracellular PLK1 protein expression via immunofluorescence assay. Compared with the cells without treatment (Blank), the expression level of PLK1 protein in HeLa cells treated with D-RNPC decreased to 44%. In contrast, no obvious inhibition was observed in cells treated with M-RNPC (Figure 4b,c). The similar inhibition trend of PLK1 protein expression in HeLa cells was also verified by Western blot (WB) analysis (Figure 4d). MTT assay showed the inhibition effect of D-RNPC to HeLa cell viability (Figure 4e). After HeLa cells were treated with D-RNPC for 72 h, the cell viability obviously decreased in comparison to the cells treated with M-RNPC. Moreover, the inhibition effect of D-RNPC was dose dependent (Figure S21, Supporting Information). In contrast, the D-RNPC did not show the inhibition effects on PLK1 protein expression and cell viability in AML-12 cells (Figure S22, Supporting Information), indicating the cell-selective protein regulation and therapy of D-RNPC.

**Figure 4.**
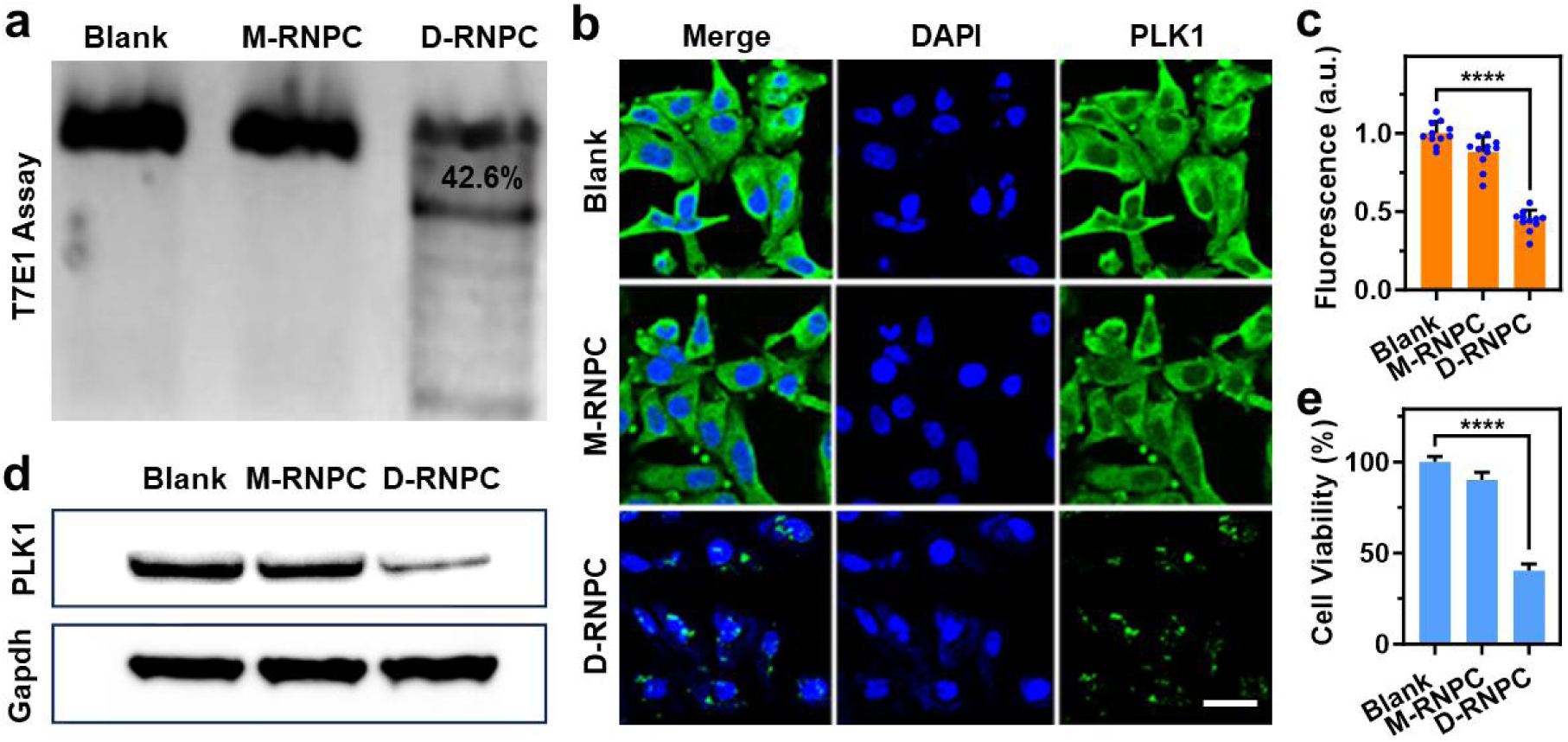
*In vitro* gene therapy of D-RNPC. a) T7E1 assay to detect genomic modifications of PLK1 in HeLa cells after incubation without (Blank) and with 200 nM D-RNPC or 200 nM M-RNPC for 72 h. Immunofluorescence analysis b) and Western blotting analysis d) of PLK1 protein in HeLa cells treated without (Blank) and with 200 nM D-RNPC or 200 nM M-RNPC for 72 h. Scale bar, 50 µm. c) Statistical analysis of PLK1 fluorescence intensity inFigure 4B. Shown are mean ± SEM from ten individual cells, ****P<0.0001. e) Cell viability of HeLa cells treated without (Blank) and with 200 nM D-RNPC or 200 nM M-RNPC for 72 h. Shown are mean ± SEM from three experiments. ****P<0.0001.

### 2.4. Construction and Performance Assessment of Folic Acid/Dual Peptide/Blocked sgRNA/Cas9 Condensates (F-D-BRNPC)

To enhance the specificity and controllability of gene editing, we designed blocked chain to hybridize with sgRNA for inhibiting the activity of RNP. In presence of survivin mRNA, the blocked chain was displaced and dissociated from sgRNA, which induced the activation of RNP for gene editing (Figure S23A, Supporting Information). We designed six blocked chains with different numbers of complementary oligonucleotides to inhibit the RNP activity. As shown in Figure S23B, Supporting Information, the blocked chain 1 induced the highest inhibitory efficiency and recovery of Cas9 activity, which was selected for D-BRNPC construction.

We next investigated the working performance of F-D-BRNPC in HeLa cells by using Cyanine 5 (Cy5) and folic acid (F) to label the blocked chain 1. Confocal microscopy images confirmed the successful construction of F-D-BRNPC (Figure S24, Supporting Information). After incubating F-D-BRNPC with HeLa cells for 12 h, the poor colocalization between RFP and Cy5 with a Pearson’s correlation coefficient of 0.37 was achieved, whereas the higher signal overlaps with Pearson’s correlation coefficients of 0.63, 0.75 and 0.62 were observed in HeLa cells treated with F-D-BRNPC in the presence of H-89 or survivin mRNA inhibitor YM155, or F-M-BRNPC assembled with blocked RNP and a misorder peptide (Figure 5a; Figure S25, Supporting Information). All these results verified that the blocked chain was released from RNP only in presence of both PKA and survivin mRNA. In addition, we also confirmed that the RNP could enter into the nucleus of HeLa cells incubated with F-D-BRNPC (Figure S26, Supporting Information). Time-dependent endocytosis of F-D-BRNPC was further investigated. As shown in Figure S27, Supporting Information, the internalization of F-D-BRNPC in HeLa cells reached saturation after 2 h incubation, which was used for the subsequent experiments. Moreover, the release of RNP was observed after 6 h incubation of F-D-BRNPC. Compared to the HeLa cells treated with D-BRNPC (D-BRNPC), the higher fluorescence signal of FAM was observed in cells incubated with F-D-BRNPC. However, the fluorescence signal was decreased upon the cell pretreated with folic acid (Figure S28, Supporting Information), indicating that the internalization of F-D-BRNPC was governed by folate receptor mediated endocytosis. Moreover, the AML-12 cells that expressed the relative low abundance of folate receptor displayed lower fluorescence signal of F-D-BRNPC compared to HeLa (Figure S29, Supporting Information), demonstrating high cell targeting and specificity of F-D-BRNPC.

**Figure 5.**
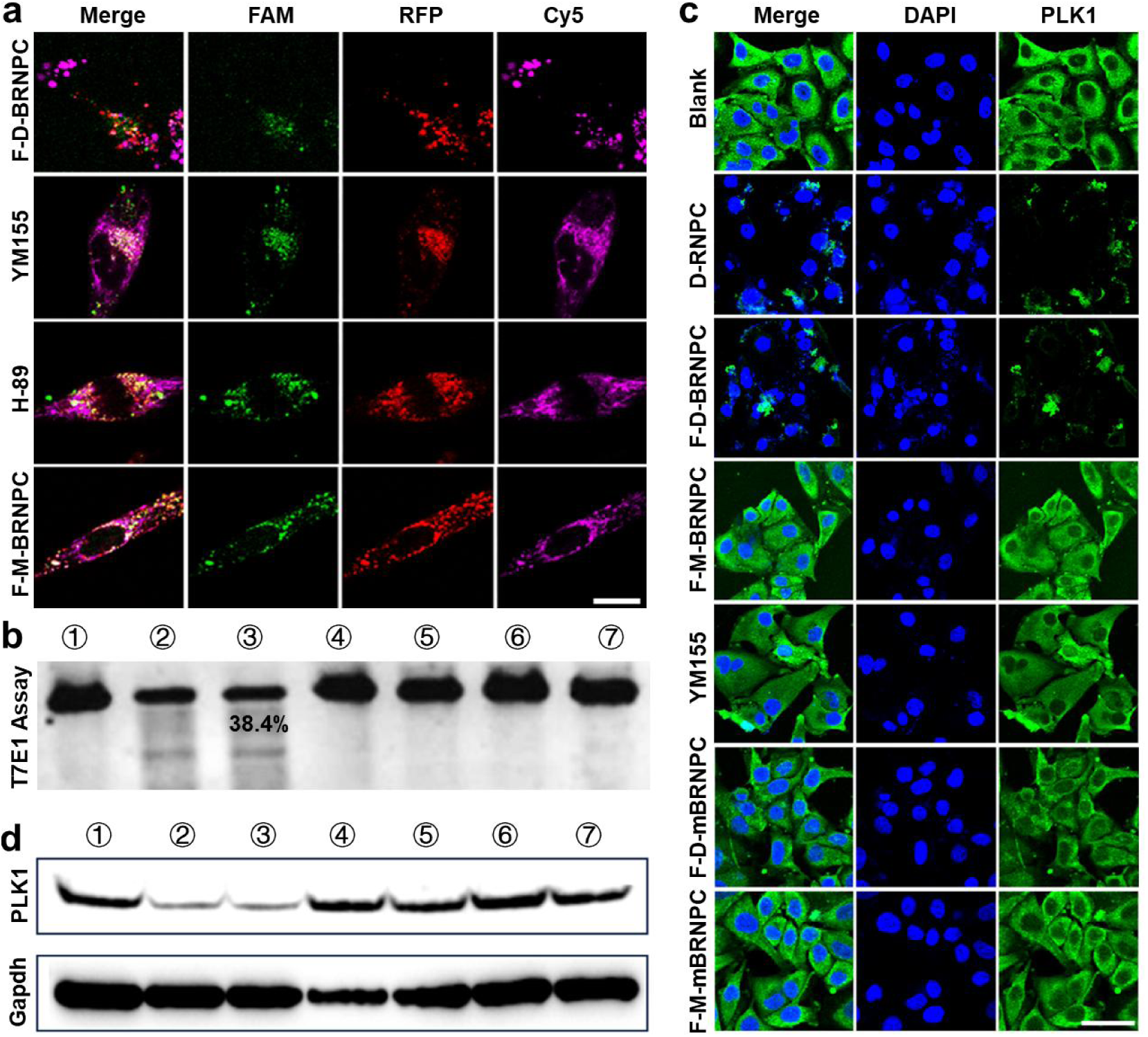
In vitro gene therapy of F-D-BRNPC. a) Confocal fluorescence imaging of HeLa cells incubated with 200 nM F-M-BRNPC, or 200 nM F-D-BRNPC in absence and presence of 20 μM H-89 (H-89) or 10 nM YM155 (YM155) for 24 h. Scale bar, 25 µm. b) T7E1 assay to detect genomic modifications of PLK1 in HeLa cells after incubation without (line 1) and with 200 nM D-RNPC (line 2), F-D-BRNPC (line 3), F-M-BRNPC (line 4), F-D-BRNPC in presence of 10 nM YM155 (line 5), F-D-mBRNPC (line 6) or F-M-mBRNPC (line7) for 72 h. c) Immunofluorescence analysis of PLK1 protein in HeLa cells treated without (Blank) and with 200 nM D-RNPC, F-D-BRNPC, F-M-BRNPC, F-D-BRNPC in presence of 10 nM YM155, F-D-mBRNPC or F-M-mBRNPC for 72 h. Scale bar, 50 µm. d) Western blotting analysis of PLK1 protein expression in HeLa cells after incubation without (line 1) and with 200 nM D-RNPC (line 2), F-D-BRNPC (line 3), F-M-BRNPC (line 4), F-D-BRNPC in presence of 10 nM YM155 (line 5), F-D-mBRNPC (line 6) or F-M-mBRNPC (line 7) for 72 h.

To further evaluate the gene editing of F-D-BRNPC in HeLa cells, we performed T7E1 assay to measure the cleavage efficiency of endogenous target *PLK1* gene. As shown in Figure 5b, the substantial target genes cleavage of approximately 38.4% was obtained in HeLa cells treated with F-D-BRNPC, similar to that in cells treated with D-RNPC. In contrast, there was no observable cleavage of PLK1 in cells incubated with F-M-BRNPC, F-D-BRNPC in presence of YM155, F-D-mBRNPC assembled with misorder blocked chain, or F-M-mBRNPC assembled with misorder blocked chain and misorder peptide. All these results testified that the gene editing of F-D-BRNPC in HeLa cells was cooperatively controlled by survivin mRNA and PKA. Immunofluorescence assay indicated that PLK1 protein expression was reduced as much as 50% by F-D-BRNPC, while negligible down-regulation of PLK1 protein was generated by F-M-BRNPC, F-D-BRNPC and YM155, F-D-mBRNPC, or F-M-mBRNPC (Figure 5c; Figure S30, Supporting Information), and the results were consistent with the WB assay (Figure 5d). MTT assay showed that F-D-BRNPC significantly reduced the viability of HeLa cells (Figure S31, Supporting Information), while no significant effect was observed in AML-12 cells (Figure S32, Supporting Information). Collectively, the results demonstrated that the high controllability and accuracy of F-D-BRNPC for gene editing, protein down-regulation and target cell therapy.

### 2.5. Formulation and Performance Characterization of Inhalable Dry Powder F-D-BRNPC

Dry powder formulation that owns greater physiochemical stability than liquid or suspension-based formulations has good aerosol performance for inhaled treatment.[34,35] The dry powder formulation of F-D-BRNPC was formed through lyophilization (Figure 6a). We subsequently tested whether the lyophilization and atomization processes could compromise the dispersity and structural integrity of the condensates. Confocal laser scanning microscopy imaging revealed no obvious changes in dispersity and morphology (Figure 6b). The mannitol is one of the commonly used excipients for improving the dispersity, stability and compatibility of dry powder formulation.[36] Upon addition of 1.5% mannitol, the yellow powder of F-D-BRNPC with excellent dispersibility was obtained after lyophilization. Moreover, the size and morphology of F-D-BRNPC were unchanged with and without mannitol (Figure S33, Supporting Information), which were similar to those of fresh liquid-based formulations. Then we assessed the stability of F-D-BRNPC dry powder. As shown in Figure S34, Supporting Information, there was no noticeable changes in the diameter of F-D-BRNPC powder after 28 days storage at 4 ℃, indicating the high stability of lyophilized F-D-BRNPC. We further evaluated the cellular internalization and gene editing of F-D-BRNPC powder in HeLa cells. The obvious fluorescence signals of RFP that largely overlapped with the DAPI fluorescence were shown in HeLa cells treated with F-D-BRNPC powder, which was consistent with the HeLa cells treated with fresh F-D-BRNPC (Figure 6c). In addition, HeLa cells incubated with F-D-BRNPC powder showed an efficient inhibition on PLK1 protein expression, similar to that in cells incubated with fresh F-D-BRNPC (Figure 6d,e,f). These results established the high stability and feasibility of F-D-BRNPC powder for cellular uptake and target gene editing.

**Figure 6.**
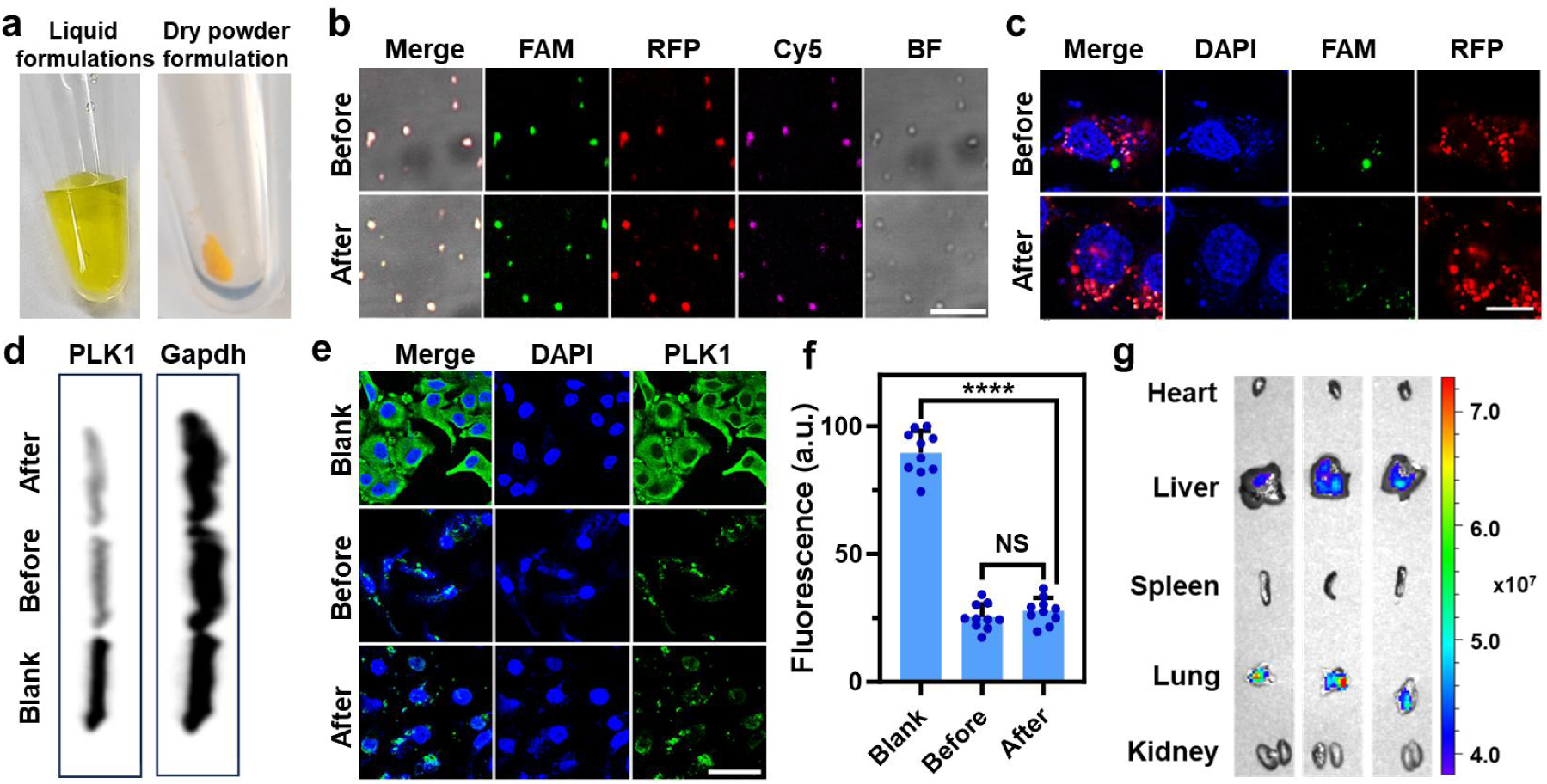
Performance characterization of inhalable dry powder F-D-BRNPC. Morphology a) confocal images of b) F-D-BRNPC before (Liquid formulation) and after (Dry powder formulation) lyophilization. Scale bar, 5 µm. c) Confocal images of HeLa cells incubated with 200 nM F-D-BRNPC condensates before (Before) and after lyophilization (After). Scale bar, 25 µm. d) Western blot analysis of PLK1 protein expression in HeLa cells incubated without (Blank) and with 200 nM F-D-BRNPC before (Before) and after lyophilization (After) for 72 h. Immunofluorescence analysis e) and Statistical analysis f) of PLK1 protein in HeLa cells incubated without (Blank) and with 200 nM F-D-BRNPC before (Before) and after lyophilization (After) for 72 h. Scale bar, 50 µm. Shown are mean ± SEM from ten individual cells, ****P<0.0001. g) *Ex vivo* images of mouse major organs that received 200 nM Cy5-labelled F-D-BRNPC after 2 h inhalation administration.

To demonstrate the feasibility of condensate for *in vivo* inhalable RNP delivery, Cy5-labelled F-D-BRNPC was used for inhaled administration to mice. *Ex vivo* imaging revealed that F-D-BRNPC are mainly enriched in the lungs after 2 h inhalation (Figure 6g; Figure S35, Supporting Information). These results indicated that the F-D-BRNPC could be used as a promising and safe carrier for inhaled delivery biomacromolecules to the lungs of mice.

### 2.6. Characterization and Application of Dual Peptide/mRNA Condensate (D-mRNAC)

To further demonstrate the versatility of enzyme-responsive condensate for biomacromolecules delivery, the mCherry mRNA was selected as a model and mixed with dual peptide for constructing dual peptide/mRNA condensate (D-mRNAC). Confocal laser scanning microscopy imaging demonstrated that the spherical D-mRNAC with high dispersibility was formed (Figure S36, Supporting Information). The optimal dual peptide concentration and incubation time for D-mRNAC assembly was 150 µM and 1 h (Figure S37, Supporting Information), respectively. After addition of 1400 nM PKA for 1 h incubation, 85% of D-mRNAC was disassembled, indicating the highly catalytic efficiency of PKA for mRNA release (Figure S38, Supporting Information).

We subsequently evaluated the feasibility of D-mRNAC for mCherry mRNA delivery and controlled protein expression in HeLa cells. As shown in Figure S39A, Supporting Information, a strong red fluorescence signal of mCherry protein was exhibited inside the cytoplasm, proving that mCherry mRNA was delivered and released inside HeLa cells. As control, no red fluorescence signal was observed in HeLa cells treated with D-mRNAC and H-89 mixture, or M-mRNAC assembled with mRNA and misorder peptide, suggesting the high specificity and controllability of D-mRNAC for mCherry mRNA release and cellular translation. In addition, almost no fluorescence of mCherry protein was shown in D-mRNAC treated AML-12 cells, indicating the cell-selective mRNA release and controlled protein expression of D-mRNAC (Figure S39B, Supporting Information). The size and morphology of D-mRNAC were almost unchanged after lyophilization, which were similar to those storage at 4 ℃ for 1 month (Figure S40, Supporting Information). Confocal laser scanning microscopy imaging certified that the intracellular performance of D-mRNAC powder was unchanged (Figure S41, Supporting Information). All these results demonstrated that the feasibility and universality of peptide-based condensate for biomacromolecules efficient delivery and controlled release.

## 3. Conclusion

In this study, we have developed a condensate based universal vehicle for biomacromolecular therapeutics controlled delivery. This delivery system has several important features, including that it could be modularly designed and simply prepared to efficiently recruit a wide range of biomacromolecules, such as CRISPR/Cas9 ribonucleoprotein (RNP) and mRNA, for formation of condensate via LLPS, leading to the activity inhibition of internal biomacromolecules due to isolation from external environment. The condensate showed high stability under various conditions, as well as lyophilization and nebulization. Moreover, the disassembly of condensate was controlled by PKA-mediated phosphorylation, which resulted in the precise release and functional recovery of loaded biomacromolecules with high efficiency and specificity.

Second, condensates have shown significant potential in drug delivery. It is noteworthy that the cellular internalization of condensates is lipid raft-mediated endocytosis, and not lysosomal-dependent endocytosis, which avoids lysosomal degradation and preserves bioactivity of cargo.[8] The condensates exhibited higher transfection efficiency and lower cytotoxicity than commercial transfection reagents of FugeneHD and Lipo2000 (Figure S19, Supporting Information). Furthermore, we displayed PKA-triggered cargo release in the cytoplasm, enabling RNP and mRNA activation for performing gene editing (Figure 4) and protein expression (Figure S39, Supporting Information), respectively. Due to PKA as a biomarker is overexpressed in cancer cell lines,[37] the enzyme-responsive condensates system could be applied for cell-selective regulation and therapy (Figure 3b; Figure S22 and S32, Supporting Information). We also modularly designed and achieved the specific cell targeting delivery, as well as endogenous PKA and RNA-triggered RNP activation (Figure S23, Supporting Information) and precise gene editing, suggesting high flexibility and programmability of the condensate for biomacromolecular therapeutics delivery and potential application in disease treatment.

The condensate with a diameter of approximately 1 μm and good storage stability in dry powder formulation is optimal for inhalable transport.[29] We have demonstrated the feasibility of condensate for aerosol inhalation delivery of RNP to lung (Figure 6). Compared to the nanoparticle and virus-based inhalable delivery systems, the condensate is more stable, biocompatible, programmable, and facile in preparation. In addition, the condensate could be modularly designed as stimuli-responsive vehicles for specific recognition of target cells and “on-demand” release of biomacromolecules induced by physicochemical triggers, which could reduce “off-target” effects and enhance therapeutic efficienc.[38,39]

In conclusion, we developed an enzyme-responsive condensate based versatile and noninvasive platform for inhaled delivery of biomolecular therapeutics, inclusive of proteins and mRNA. We envision that the modular, stable, highly biocompatible and controllable vehicle system can be broadly applied to drug delivery and cell regulation, potentially opening new avenues for diseases treatment and life science research.

## 4. Experimental Section/Methods

*Reagents*: Unless otherwise specified, all chemicals were purchased from Sigma. Use commercial reagents without further purification. All aqueous solutions were prepared using ultrapure water (18.2 MΩ·cm, Milli-Q, Millipore). LysoTracker blue, 4’,6-Diamidino-2-Phenylindole (DAPI), bovine serum albumin Fetal bovine serum (FBS), agarose, 40% polyacrylamide, TEMED, APS and all of the DNA were from Sangon Biotech Co., Ltd (Shanghai, China). Protein kinase A catalytic subunit from bovine heart from Sigma-Aldrich Trading Co., Ltd (Shanghai, China). T7 endonuclease I was purchased from New England Biolabs, Ltd (Beijing, China). Anti-PLK1 antibody (ab17056) was obtained from Abcam (Shanghai, China). PLK1 polyclonal antibody was purchased from Qiyan Biotechnology Co., Ltd (Beijing, China). GAPDH antibody (HRP conjugated) was obtained from Shenger Biotechnology Co., Ltd (Shanghai, China). ECL plus ultra sensitive luminescent liquid, DNA extraction kit was from Yifei Xue Biotechnology Co., Ltd (Nanjing, China). Universal DNA Purification and Recovery Kit from Yuanye Bio-Technology Co., Ltd (Shanghai, China), MTT Cell Proliferation and Cytotoxicity Detection Kit from Beyotime Biotechnology Co., Ltd (Shanghai, China), 2×Taq PCR Master Mix and Horseradish enzyme labeled goat anti mouse IgG (H+L) was from Yizhi Mei Biotechnology Co., Ltd (Nanjing, China). Alt-RS.p.Cas9 (RFP) was purchased from Integrated DNA Technologies, Inc. (Iowa, US). sgRNA was purchased from GenScript Biotechnology Co., Ltd (Nanjing, China). All oligonucleotides are listed in Supplemental information, Table S1.

*Apparatus:* All the intracellular images were taken by a Nikon A1& SIM-S&STORM super resolution microscope (Tokyo, Japan). All the *in vitro* fluorescence measurements were measured on an F-7000 spectrometer (HITACHI, Japan). The concentrations of nucleic acids were measured using a NanoDrop One UV-Vis spectrophotometer. The gel electrophoresis was performed on a Tanon EPS-300 Electrophoresis Analyser (Tanon Science & Technology Company, China) and imaged on Bio-rad ChemDoc XRS (Bio-Rad, USA). The zeta potential and dynamic lights scattering (DLS) were measured on a Zetasizer Nano ZS90 device (Malvern, UK). MTT assays were measured with a Safire microplate Analyzer (Molecular Devices, America).

*Construction of peptide/RNP condensates (RNPC):* The red fluorescent protein (RFP) fused Cas9 and sgRNA were mixed at 1:1 molar ratio in a buffer consisting of 20 mM HEPES, 2.5 mM ATP, and 1.6 mM dithiothreitol, pH 7.4. The mixture was incubated at 37 °C for 10 min to facilitate the formation of the Cas9/sgRNA ribonucleoprotein complex (RNP). Subsequently, the RNP was mixed with peptides for 1 h to obtain RNPC. Specifically, for D-RNPC, T-RNPC and M-RNPC formation, the final concentrations of dual repeat peptide, triple repeat peptide and misordered peptide were 200 µM, 75 µM, and 200 µM, respectively, and the final concentration of RNP was 200 nM. We use the concentration of RNP as the final concentration of RNPC.

For F-D-BRNPC (or F-D-mBRNPC) construction, the folic acid block chain 1 (or misorder block chain 1) was mixed with sgRNA at 1:1 molar ratio and incubated at 90 °C for 15 min. Then, the F-D-BRNPC (or F-D-mBRNPC) was prepared by using of blocked sgRNA and following the above procedure.

For peptide concentration optimization, 200 nM RNP was mixed with different concentration of dual-peptide or triple peptide.

For sgRNA/Cas9 ratio optimization, sgRNA was mixed with Cas9 at different molar ratios (sgRNA: Cas9 = 1:2, 1:1, 2:1, 3:1, 4:1), and subsequently the dual-peptide and triple peptide at optimal concentration were added to form D-RNPC and T-RNPC, respectively. The final concentration of sgRNA was 200 nM.

For *in vitro* RNPC imaging, 10 μL of the RNPC was added onto the confocal dish and visualized by using a laser scanning confocal microscope (Nikon A1& SIM-S). Images were collected with an excitation at 488 nm (FAM) and 532 nm (RFP) using a 100 × oil immersion objective. Image analysis was performed using Image J software.

*PKA triggered RNPC decomposition:* The 200 nM D-RNPC, T-RNPC, and M-RNPC were prepared under optimal conditions. RNPC was mixed with and without PKA at a final concentration of 1.4 µM for 1 h, then was imaged with Nikon A1& SIM-S&STORM super-resolution microscope. Images were collected with an excitation at 488 nm (FAM) using a 100 × oil immersion objective. Image analysis was performed using Image J software.

*Fluorescence assay of RNPC formation and decomposition:* For fluorescence assay of RNPC formation, FAM-5 labeled peptides (200 µM dual repeat peptide, 75 µM triple repeat peptide and 200 µM misordered peptide) were mixed with 200 nM RNP in a buffer solution containing 20 mM HEPES, 2.5 mM ATP, and 1.6 mM dithiothreitol (pH 7.4). The supernatant of mixture was collected via centrifugation after incubation for different times (0 min, 10 min, 20 min, 30 min, 60 min, and 120 min). Then the fluorescence of the supernatant was measured with excitation at 488 nm, and emission at 518 nm.

To measure the dynamic of PKA-triggered RNPC decomposition, 200 nM D-RNPC, T-RNPC or M-RNPC was mixed with PKA at final concentration of 1.4 µM. After incubation for different times (0 min, 10 min, 20 min, 40 min, 90 min, and 180 min), the supernatant of mixture was collected via centrifugation, and the fluorescence of the supernatant was measured with excitation at 488 nm, and emission at 518 nm.

*RNP activity characterization: PLK1* gene (500 ng) was incubated with 200 nM RNP in NEB Buffer 2 for 1 h at 37 ℃. Then, the mixture was analyzed by 2% agarose gel electrophoresis.

*Cell culture:* Human cervix carcinoma (HeLa) cells lines, Alpha Mouse Liver 12 (AML-12) cells lines (Procell Life Science & Technology, Wuhan, China) were cultured in Dulbecco’s modified Eagle’s medium (DMEM) supplemented with 10% fetal bovine serum (FBS), 100 mg/mL streptomycin and 100 U/mL penicillin-streptomycin at 37 ℃ in a humidified incubator containing 5% CO2 and 95% air. Short tandem repeats (STR) profiling and mycoplasma testing were conducted for each cell line before use. Cell numbers were determined with a Petroff-Hausser cell counter (USA).

*Cellular uptake of D-RNPC:* To optimize the cellular uptake times of D-RNPC, HeLa cells were seeded in confocal culture dishes at a density of 2×10^3^ cells per well and incubated with 200 nM D-RNPC for different times (6 h, 12 h, 24 h, 48 h, and 72 h). Then, the cells were washed three times with PBS and incubated with 1 μg/mL 4’,6-diamidino-2-phenylindole (DAPI) for 15 min. Subsequently, the cells were washed three times with PBS and imaged with the Nikon A1& SIM-S&STORM super-resolution microscope (Tokyo, Japan). The intracellular fluorescence intensity was measured using ImageJ. FAM and DAPI were excited with 488 nm and 405 nm lasers, respectively.

*Cell uptake mechanism assay:* HeLa cells were seeded in confocal culture dishes at a density of 2×10^3^ cells per well, and incubated with inhibitors of chlorpromazine (30 μM), amiloride (20 μM), methyl-β-cyclodextrin (2.5 mM), or sodium azide (100 mM) for 30 min. Subsequently, 200 nM D-RNPC was added and incubated for 12 h. Then, the cell was incubated with 1 μg/mL DAPI for 15 min before imaging. RFP, FAM and DAPI were excited with 532 nm, 488 nm and 405 nm lasers, respectively.

*Cell co-localization assay:* HeLa cells were seeded in confocal culture dishes at a density of 2×10^3^ cells per well, then incubated them with 200 nM D-RNPC for 24 h. The cells were then stained with Lyso Tracker Blue for 30 min. Afterward, the cells were washed three times with PBS before imaging. RFP, FAM and DAPI were excited with 532 nm, 488 nm and 405 nm lasers, respectively.

*Immunofluorescence analysis:* HeLa cells were seeded in confocal culture dish at a density of 2×10^3^ cells per well for 24 h at 37 °C. Then, cells were incubated without and with 200 nM M-RNPC or D-RNPC in absence and presence of 20 μM PKA inhibitor N-[2-(p-Bromocinnamylimino)ethyl]-5-isoquinolinesulfonamide·2HCl hydrate (H-89) for 72 h. Then, the cells were fixed by 4% paraformaldehyde for 10 min and permeabilized with 0.2% Triton X-100 for 15 min. Subsequently, the cells were pre-blocked in PBS containing 10% fetal bovine serum (v/v) and 5% bovine serum albumin (w/v) for 1 h, and incubated with PLK1 antibody at room temperature for 2 h. After incubation with FITC goat Anti-Rabbit IgG for 1 h at room temperature, the cells were stained with 1 μg/mL DAPI for 15 min and observed under Nikon A1& SIM-S&STORM super resolution microscope. FITC and DAPI were excited with 488 nm and 405 nm lasers, respectively.

*Western blotting analysis:* HeLa cells were seeded into six well plates to a density of 2 × 10^5^ per well for 24 h. Then the cells were incubated without and with 200 nM M-RNPC or D-RNPC for 72 h. The cells were washed by ice-cold PBS and treated with 200 μL SDS lysis buffer. The protein concentration was accessed by BCA protein quantification Kit, separated by 10% SDS-PAGE and transferred to PVDF membranes. Then membranes were blocked with 5% skimmed milk at room temperature for 2 h, followed by incubating with 5% BSA in which anti-PLK1 antibody was contained at 4 °C overnight. Afterwards, the membranes were incubated with 5% skimmed milk in which peroxidase affinipure goat anti-mouse IgG were contained at room temperature for 1 h. Super ECL detection reagent was used for signal detection.

*T7E1 assay:* HeLa cells were seeded six well plate to a density of 2 × 10^5^ per well for 24 h. Then the cells were incubated without and with 200 nM M-RNPC or D-RNPC for 72 h. Subsequently, the cells were washed three times with PBS, and the total genomic DNA was extracted by using a DNA extraction kit. After amplification the target gene with PCR, the product was cleaved with 1 uL T7 endonuclease 1 for 30 min, and analyzed with 2% agarose gel electrophoresis. The cutting percentage was analyzed using ImageJ statistical software.

*3-(4,5-dimethylthiazol-2-yl)-2,5-diphenyltetrazolium bromide (MTT) assay:* HeLa (or AML-12) cells were cultured at a density of 1×10^5^ cells per chamber in a 96-well plate for 24 h. After removing the culture medium, the cells were washed with PBS and incubated with different concentrations of D-RNPC (0 nM, 1 nM, 10 nM, 50 nM, 100 nM, and 200 nM) for an additional 72 h. Following this incubation, the cells were washed twice with washing buffer. Next, 50 μL of 5 mg/mL tetrazolium salt solution was added, and the cells were incubated for 4 h. After removing remaining tetrazolium salt solution, 100 μL of dimethyl sulfoxide (DMSO) was added, and the plate was shaken on a shaker for 10 min. Finally, the optical density was measured at a wavelength of 490 nm using a SAFIRE microplate analyzer.

*Intracellular performance of F-D-BRNPC:* HeLa cells were seeded in confocal culture dish at a density of 2×10^3^ cells per well for 24 h at 37 °C. Then the cells were treated without and with 200 nM F-M-RNPC, or F-D-BRNPC in presence and absence H-89 (20 μM) or survivin mRNA inhibitor YM155 (10 nM) for 12 h. After washing three times with PBS, the cells were incubated with 1 μg/mL DAPI for 15 min before imaging. RFP, FAM and Cy5 were excited with 532 nm, 488 nm and 650 nm lasers, respectively.

*Construction of peptide / mRNA condensates (D-mRNAC):* 200 nM mCherry mRNA was mixed with 150 µM dual peptides in a buffer consisting of 20 mM HEPES, 2.5 mM ATP, and 1.6 mM dithiothreitol at pH 7.4. The mixture was incubated at 37 °C for 1 h to obtain D-mRNAC. We use the concentration of mRNA as the final concentration of D-mRNAC.

*Cellular uptake of D-mRNAC:* HeLa cells were seeded in confocal culture dishes at a density of 2×103 cells per well. Then the cells were co-cultured with 200 nM mCherry mRNA, 150 µM dual peptides, 200 nM D-mRNAC, 200 nM D-mRNAC (20 μM H-89 incubation cells for 30 min), 200 nM M-mRNAC for 24 h. Then, the cells were washed three times with PBS. Imaged with the Nikon A1& SIM-S&STORM super-resolution microscope (Tokyo, Japan). FAM and mCherry were excited with 488 nm and 587 nm lasers, respectively. *Animals:* All animal studies were conducted at Nanjing University of Science and Technology and in compliance with the regulations of the Ethics Committee for the Care and Use of Laboratory Animals. The animal experiment was approved by the Institutional Review Committee of Nanjing University of Science and Technology (protocol code: ACUC-NUST-20250228013).

*F-D-BRNPC inhaled in vivo delivery:* Female C57BL6 mice aged 5 to 6 weeks were loaded with an equal dose of F-D-BRNPC solution into a nebulizer, and air flow was used to guide the aerosol along the gasket into the chamber until no more aerosol was observed. Mice treated with PBS were used as negative controls. Mice were euthanized 2 h after nebulization, and the major organs (heart, liver, spleen, Lung, and Kidney) were harvested and imaged on an IVIS system (PerkinElmer, USA). All animal experiments have been carried out according to regulations and guidelines on animal use from the Animal Care and Use Committee of Nanjing University of Science and Technology.

*Statistical analysis:* All grayscale, colocalization, and fluorescence intensity analyses were conducted using ImageJ. Statistical analysis was performed using GraphPad Prism 7.0, and all data were expressed as mean ± standard deviation. For two groups, a student’s t-test was conducted, while analysis of variance (ANOVA) was performed for multiple groups. When P < 0.05, the difference between the control group is considered significantly.

## Supporting Information

Supporting Information is available from the Wiley Online Library or from the author.

## Acknowledgements

The authors gratefully acknowledge the National Natural Science Foundation of China (22174066, 22104058, 22374076), the Natural Science Foundation of Jiangsu Province (BK20200459, BK20231455), the Program of Jiangsu Specially-Appointed Professor, Fundamental Research Funds for the Central Universities (30922010501, 30924010809). State Key Laboratory of Analytical Chemistry for Life Science (SKLACLS2412).

## Author contributions

Y.X.: investigation, data curation, writing-original draft, methodology; S.J.Z., T.Z., Y.L.D., X.Y.G., Q.Y.H., and L.W.: investigation; S.Y.D.: conceptualization, supervision, validation; K.W.R.: supervision, conceptualization, project administration, writing – review & editing. All authors discussed the results and reviewed the manuscript.

## Data Availability Statement

The authors declare that the data supporting the findings of this study are available within the manuscript or supplemental information. Raw data will be made available from the lead contact upon reasonable request.

Received: ((will be filled in by the editorial staff)) Revised: ((will be filled in by the editorial staff)) Published online: ((will be filled in by the editorial staff))

---

**ToC Figure.**
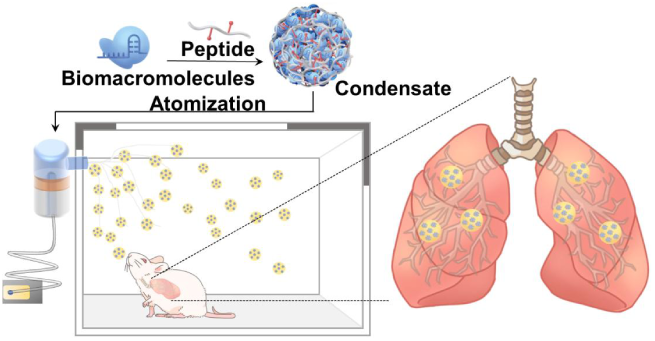
A highly biocompatible and controllable platform based on enzyme-responsive condensates for inhalation delivery of biomolecules. Modular design can be used to rapidly recruit and enrich biomolecules, including Cas9 ribonucleoprotein (RNP) and mRNA into condensates. After nebulization administration, the condensates mediate the delivery of biomolecules to the lungs.

## References

[1] M. P. Stewart, A. Sharei, X. Ding, G. Sahay, R. Langer, K. F. Jensen, Nature 2016, 538, 183–192.

[2] S. F. Dowdy, Nat. Biotechnol. 2017, 35, 222–229.

[3] D. M. Rad, M. A. Rad, S. R. Bazaz, N. Kashaninejad, D. Jin, M. Ebrahimi Warkiani, Adv. Mater. 2021, 33, 2005363.

[4] X. Sun, S. Setrerrahmane, C. Li, J. Hu, H. Xu, Sig. Transduct. Target. Ther. 2024, 9, 316.

[5] C. Liu, T. Wan, H. Wang, S. Zhang, Y. Ping, Y. Cheng, Sci. Adv. 2019, 5, eaaw8922.

[6] J. V. Haasteren, J. Li, O. J. Scheideler, N. Murthy, D. V. Schaffer, Nat. Biotechnol. 2020, 38, 845–855.

[7] W. Tai, P. Zhao, X. Gao, Sci. Adv. 2020, 6, eabb0310.

[8] Y. Sun, S. Y. Lau, Z. W. Lim, S. C. Chang, F. Ghadessy, A. Partridge, A. Miserez, Nat. Chem. 2022, 14, 274–283.

[9] D. Wang, P. W. L. Tai, G. Gao, Nat. Rev. Drug Discov. 2019, 18, 358–378.

[10] L. Ringgaard, F. Melander, R. Eliasen, J. R. Henriksen, R. I. Jølck, T. B. Engel, M. Bak, F. P. Fliedner, K. Kristensen, D. R. Elema, A. Kjaer, A. E. Hansen, T. L. Andresen, Sci. Adv. 2020, 6, eaba5628.

[11] X. Han, M.-G. Alameh, N. Gong, L. Xue, M. Ghattas, G. Bojja, J. Xu, G. Zhao, C. C. Warzecha, M. S. Padilla, R. El-Mayta, G. Dwivedi, Y. Xu, A. E. Vaughan, J. M. Wilson, D. Weissman, M. J. Mitchell, Nat. Chem. 2024, 16, 1687–1697.

[12] E. Tan, T. Wan, Q. Pan, J. Duan, S. Zhang, R. Wang, P. Gao, J. Lv, H. Wang, D. Li, Y. Ping, Y. Cheng, Sci. Adv. 2024, 10, eadl4336.

[13] A. Suberi, M. K. Grun, T. Mao, B. Israelow, M. Reschke, J. Grundler, L. Akhtar, T. Lee, K. Shin, A. S. Piotrowski-Daspit, R. J. Homer, A. Iwasaki, H.-W. Suh, W. M. Saltzman, Sci. Transl. Med. 2023, 15, eabq0603.

[14] L. Rotolo, D. Vanover, N. C. Bruno, H. E. Peck, C. Zurla, J. Murray, R. K. Noel, L. O’Farrell, M. Araínga, N. Orr-Burks, J. Y. Joo, L. C. S. Chaves, Y. Jung, J. Beyersdorf, S. Gumber, R. Guerrero-Ferreira, S. Cornejo, M. Thoresen, A. K. Olivier, K. M. Kuo, J. C. Gumbart, A. R. Woolums, F. Villinger, E. R. Lafontaine, R. J. Hogan, M. G. Finn, P. J. Santangelo, Nat. Mater. 2023, 22, 369–379.

[15] Y. Zhang, T. Sun, C. Jiang, Acta Pharm. Sin. B. 2018, 8, 34–50.

[16] A. Kakkar, G. Traverso, O. C. Farokhzad, R. Weissleder, R. Langer, Nat. Rev. Chem. 2017, 1, 0063.

[17] R. Goswami, T. Jeon, H. Nagaraj, S. Zhai, V. M. Rotello, Trends Pharmacol. Sci. 2020, 41, 743–754.

[18] S. Yu, H. Yang, T. Li, H. Pan, S. Ren, G. Luo, J. Jiang, L. Yu, B. Chen, Y. Zhang, S. Wang, R. Tian, T. Zhang, S. Zhang, Y. Chen, Q. Yuan, S. Ge, J. Zhang, N. Xia, Nat. Commun. 2021, 12, 5131.

[19] S. Alberti, A. Gladfelter, T. Mittag, Cell 2019, 176, 419–434.

[20] S. F. Banani, H. O. Lee, A. A. Hyman, M. K. Rosen, Nat. Rev. Mol. Cell Biol. 2017, 18, 285–298.

[21] Y. Sun, X. Wu, J. Li, M. Radiom, R. Mezzenga, C. S. Verma, J. Yu, A. Miserez, Nat. Commun. 2024, 15, 10094.

[22] P. Wen, H. Huang, R. Zhang, H. Zheng, T. Liang, C. Zhuang, Q. Wu, J. Wang, F. Liu, K. Zhang, W. Wu, K. He, F. Liu, H. Li, Z. Gu, Nat. Chem. 2025, 17, 279–288.

[23] M. Miragoli, P. Ceriotti, M. Iafisco, M. Vacchiano, N. Salvarani, A. Alogna, P. Carullo, G. B. Ramirez-Rodríguez, T. Patrício, L. D. Esposti, F. Rossi, F. Ravanetti, S. Pinelli, R. Alinovi, M. Erreni, S. Rossi, G. Condorelli, H. Post, A. Tampieri, D. Catalucci, Sci. Transl. Med. 2018, 10, eaan6205.

24. A. K. D. Popowski, A. Moatti, G. Scull, D. Silkstone, H. Lutz, B. López de Juan Abad, A. George, E. Belcher, D. Zhu, X. Mei, X. Cheng, M. Cislo, A. Ghodsi, Y. Cai, K. Huang, J. Li, A. C. Brown, A. Greenbaum, P.-U. C. Dinh, K. Cheng, Matter 2022, 5, 2960–2974.

[24] X. Bai, Q. Chen, F. Li, Y. Teng, M. Tang, J. Huang, X. Xu, X.-Q. Zhang, Nat. Commun. 2024, 15, 6844.

[25] C. Shaffer, Nat. Biotechnol. 2020, 38, 1110–1112.

[26] Z. Li, Z. Wang, P.-U. C. Dinh, D. Zhu, K. D. Popowski, H. Lutz, S. Hu, M. G. Lewis, A. Cook, H. Andersen, J. Greenhouse, L. Pessaint, L. J. Lobo, K. Cheng, Nat. Nanotechnol. 2021, 16, 942–951.

[27] S. Liu, Y. Wen, X. Shan, X. Ma, C. Yang, X. Cheng, Y. Zhao, J. Li, S. Mi, H. Huo, W. Li, Z. Jiang, Y. Li, J. Lin, L. Miao, X. Lu, Nat. Commun. 2024, 15, 9471.

[28] M. Liu, S. Hu, N. Yan, K. D. Popowski, K. Cheng, Nat. Nanotechnol. 2024, 19, 565–575.

[29] S. Gou, G. Wang, Y. Zou, W. Geng, T. He, Z. Qin, L. Che, Q. Feng, K. Cai, Adv. Mater. 2023, 35, 2303718.

[30] W. M. Aumiller, C. D. Keating, Nat. Chem. 2016, 8, 129–137.

[31] N. Song, Y. Chu, S. Li, Y. Dong, X. Fan, J. Tang, Y. Guo, G. Teng, C. Yao, D. Yang, Sci. Adv. 2023, 9, eadi3602.

[32] P. Panja, N. R. Jana, J. Phys. Chem. B. 2020, 124, 5323–5333.

[33] T. Parumasivam, R. Y. K. Chang, S. Abdelghany, T. T. Ye, W. J. Britton, H.-K. Chan, Adv. Drug Deliver. Rev. 2016, 102, 83–101.

[34] Y. Ye, Y. Ma, J. Zhu, Int. J. Pharmaceut. 2022, 614, 121457.

[35] S. D. Anderson, E. Daviskas, J. D. Brannan, H. K. Chan, Adv. Drug Deliver. Rev. 2018, 133, 45–56.

[36] Y. S. Cho, Y. G. Park, Y. N. Lee, M.-K. Kim, S. Bates, L. Tan, Y. S. Cho-Chung, Proc. Natl. Acad. Sci. U.S.A. 2000, 97, 835–840.

[37] J. Kim, Y. Eygeris, R. C. Ryals, A. Jozić, G. Sahay, Nat. Nanotechnol. 2024, 19, 428–447.

[38] S. K. Alsaiari, B. Eshaghi, B. Du, M. Kanelli, G. Li, X. Wu, L. Zhang, M. Chaddah, A. Lau, X. Yang, R. Langer, A. Jaklenec, Nat. Rev. Mater. 2025, 10, 44–61.

