## Supporting Information for "Phase-separating enzyme-responsive condensates for inhalable delivery of biomacromolecules"

Nanjing 210094 (China)

Y. Du, X. Guo, Prof. S. Deng

School of Environmental and Biological Engineering

Nanjing University of Science and Technology

Nanjing 210094 (China)

This Supplemental file includes:

Figure S1-S41

Tables S1

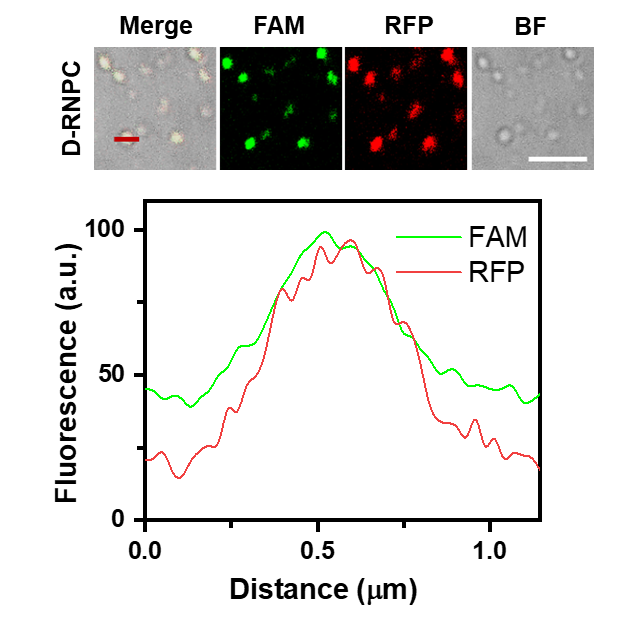

**Figure S1.** Confocal images and fluorescence intensity analysis of D-RNPC condensates. Scale bar, 5 µm.

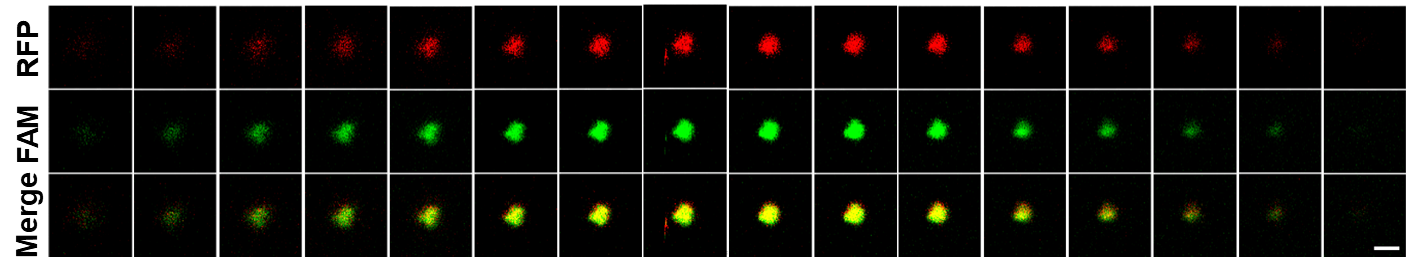

**Figure S2.** Z-stack images of D-RNPC condensates. 0.1 microns per layer. Scale bar, 2 µm.

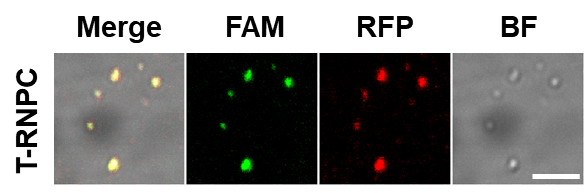

**Figure S3.** Confocal images of T-RNPC condensates. Scale bar, 5 µm.

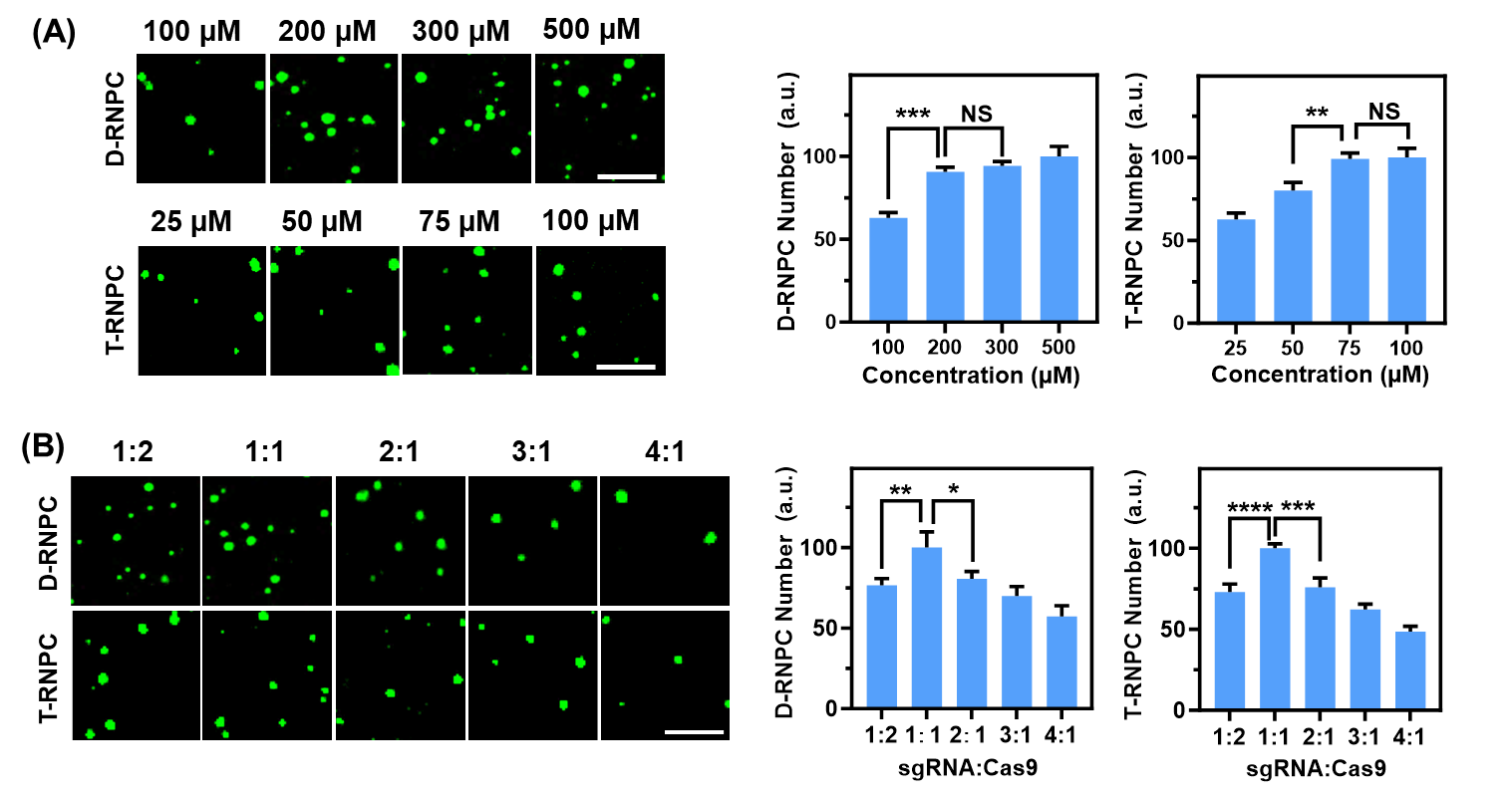

**Figure S4.** Confocal images and number statistics of D-RNPC and T-RNPC. (**A**) Effect of peptide concentrations on D-RNPC and T-RNPC preparation. Scale bar, 5 µm. Shown are the mean and the standard error of the mean (SEM) values from at least three representative images. Two-tailed Student's t-test: **P<0.01, ***P<0.001. (**B**) Effect of the ratio of sgRNA to Cas9 on D-RNPC and T-RNPC preparation. Scale bar, 5 µm. Shown are the mean and the standard error of the mean (SEM) values from at least three representative images. Two-tailed Student's t-test: *P<0.1, **P<0.01, ***P<0.001, ****P<0.0001.

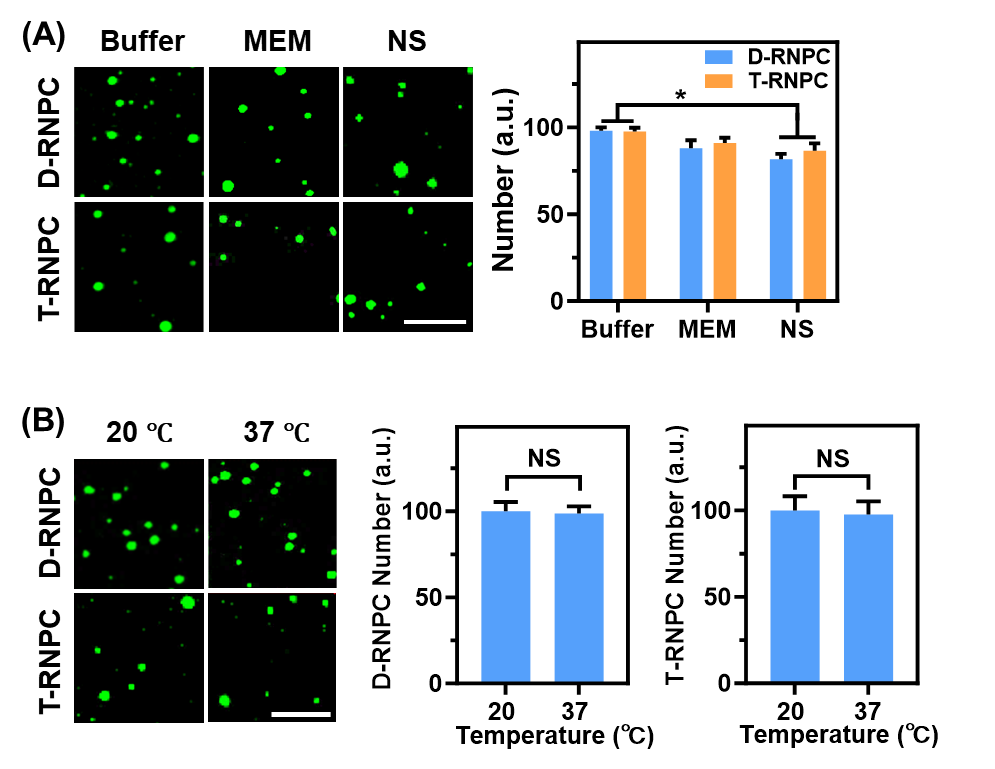

**Figure S5.** The stability assay of D-RNPC and T-RNPC. (**A**) The stability of D-RNPC and T-RNPC in solution of Buffer (20 mM HEPES, 2.5 mM ATP, and 1.6 mM dithiothreitol, pH 7.4), minimal essential medium (MEM) and normal saline (NS). Scale bar, 5 µm. Shown are the mean and the standard error of the mean (SEM) values from at least three representative images. Two-tailed Student's t-test: *P < 0.1. (**B**) The stability of D-RNPC and T-RNPC under 37 ℃ and 20 ℃. Shown are the mean and the standard error of the mean (SEM) values from at least three representative images..

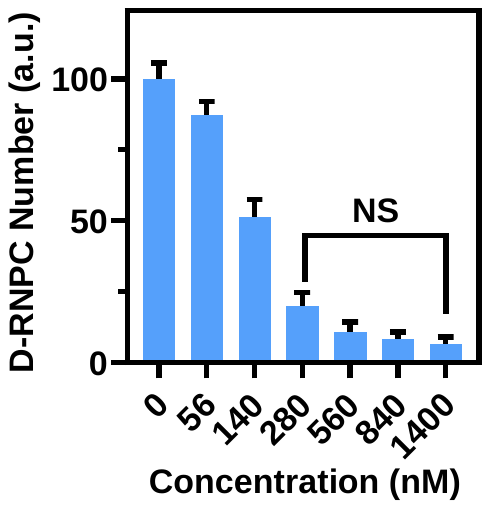

**Figure S6.** PKA triggered D-RNPC decomposition. The number statistics of D-RNPC after incubation with different concentration of PKA for 60 min. Shown are the mean and the standard error of the mean (SEM) values from at least three representative images.

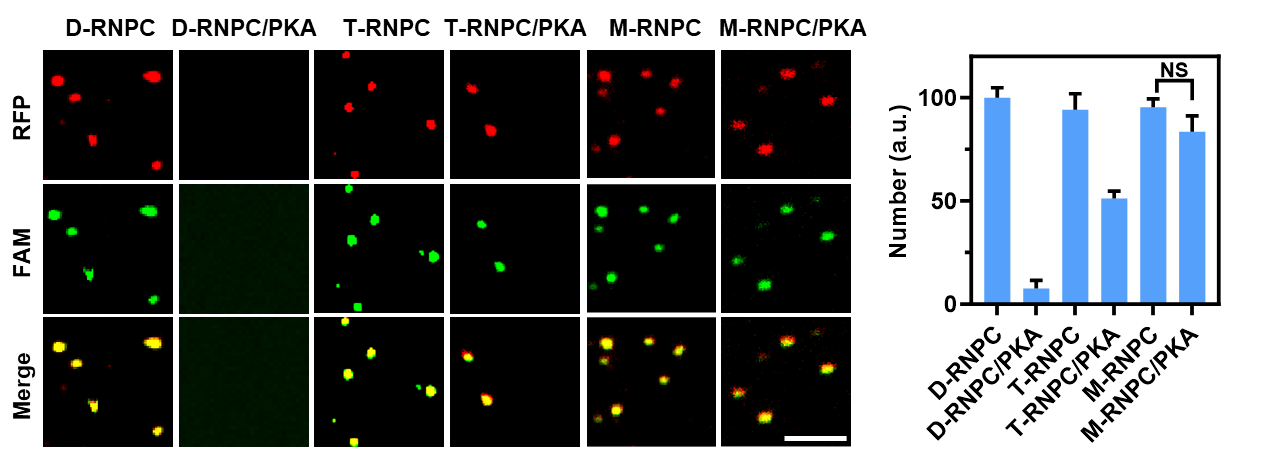

**Figure S7.** Response of different RNP condensates to PKA. Confocal images and number statistics of 200 nM D-RNPC, T-RNPC, or M-RNPC before and after incubation with PKA (1400 nM) for 1 h. Scale bar, 5 µm. Shown are the mean and the standard error of the mean (SEM) values from at least three representative images.

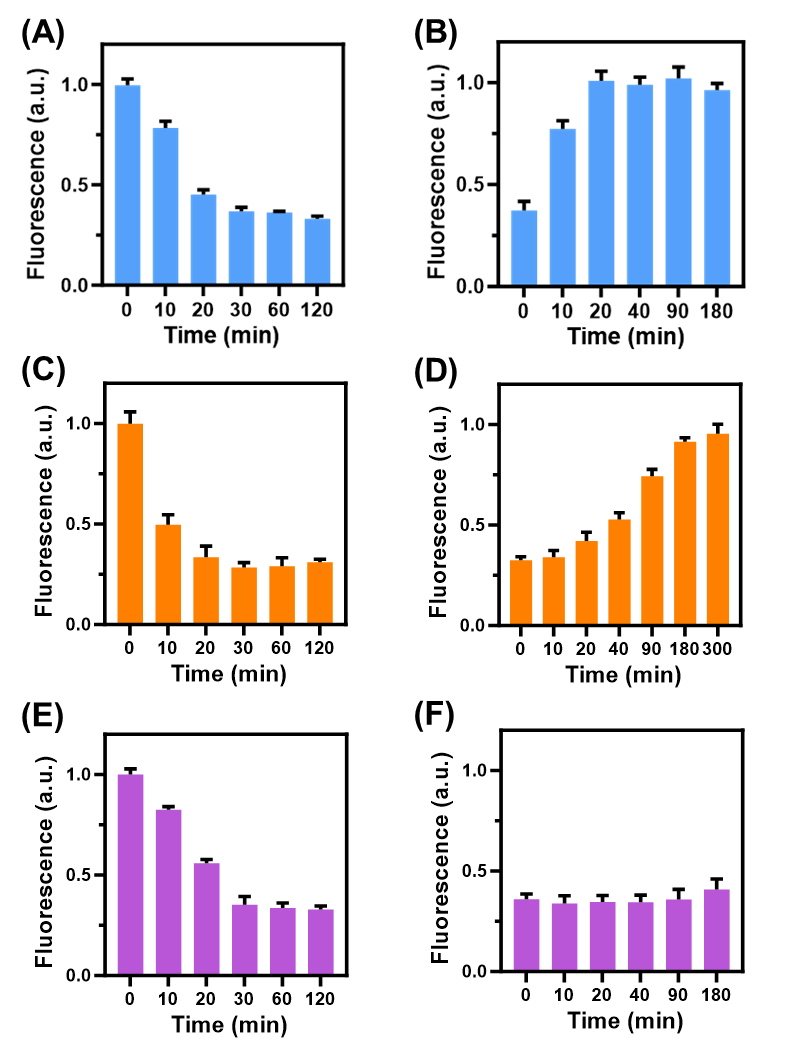

**Figure S8.** Fluorescence analysis formation and decomposition of different RNP condensates. Fluorescence intensity of the supernatant after incubation of peptide and RNP for different times in 200 nM D-RNPC (**A**), 200 nM T-RNPC (**C**) and (**E**) 200 nM M-RNPC. Fluorescence intensity of the supernatant in 200 nM D-RNPC (**B**), 200 nM T-RNPC (**D**) and (**F**) 200 nM M-RNPC upon incubation with 1400 nM PKA for different times. Shown are the mean and the standard error of the mean (SEM) values from at least three representative images.

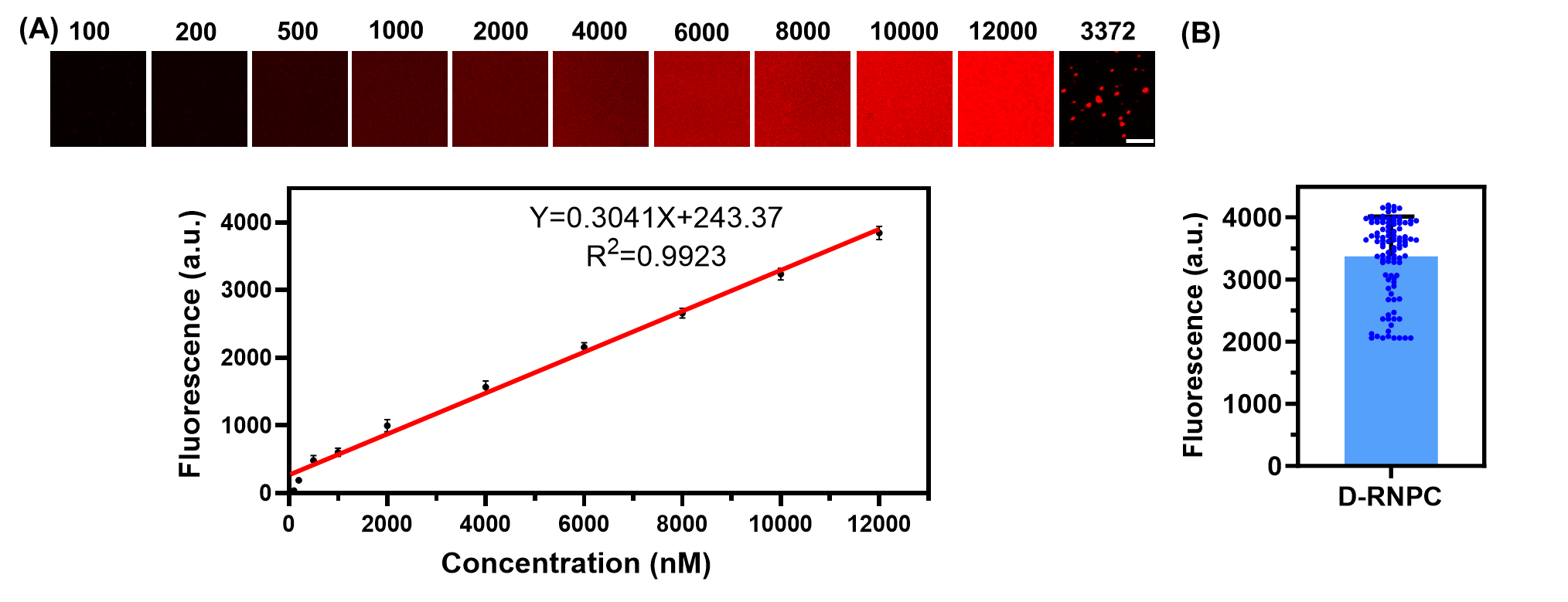

**Figure S9.** Cas9 enrichment in D-RNPC. (**A**) Imaging and the correlation between Cas9 concentration (100-12000 nM) and fluorescence signal intensities. Scale bar, 5 µm. Shown are mean ± SEM (n = 3). (**B**) The fluorescence of D-RNPC shown in panel A. Each data point represents one condensate. Shown are the mean and SEM values from three representative images.

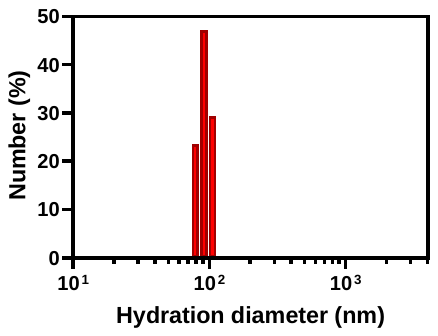

**Figure S10.** DLS analysis of hydrodynamic diameter distribution of RNP.

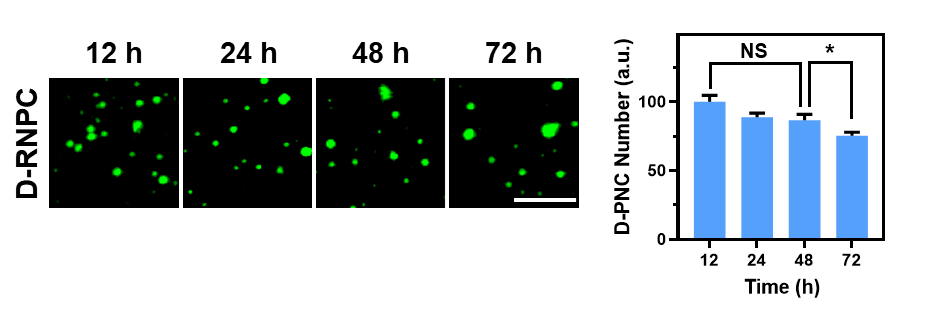

**Figure S11.** Confocal images and number statistics of D-RNPC at different incubation time. Scale bar, 5 µm. Shown are the mean and the standard error of the mean (SEM) values from at least three representative images. Two-tailed Student's t-test: *P < 0.1.

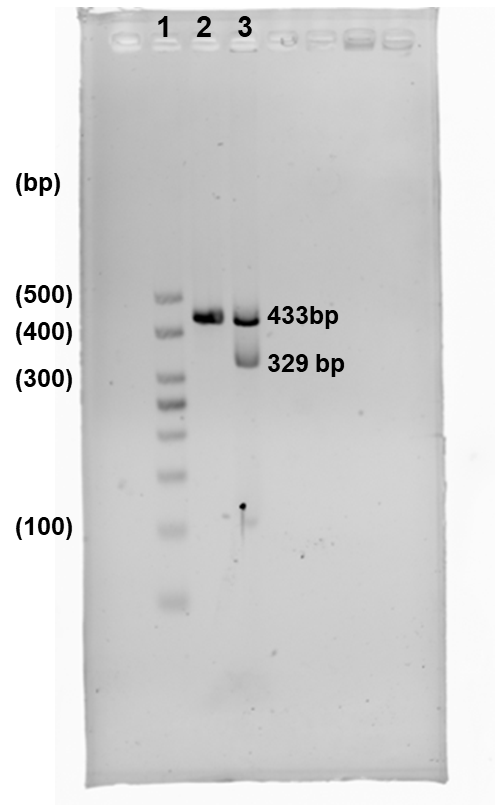

**Figure S12.** Gene editing efficiency of RNP. The 2% agarose gel electrophoresis analysis gene editing efficiency of RNP. 1. DNA Ladder, 2. Target DNA, 3. Target DNA + RNP.

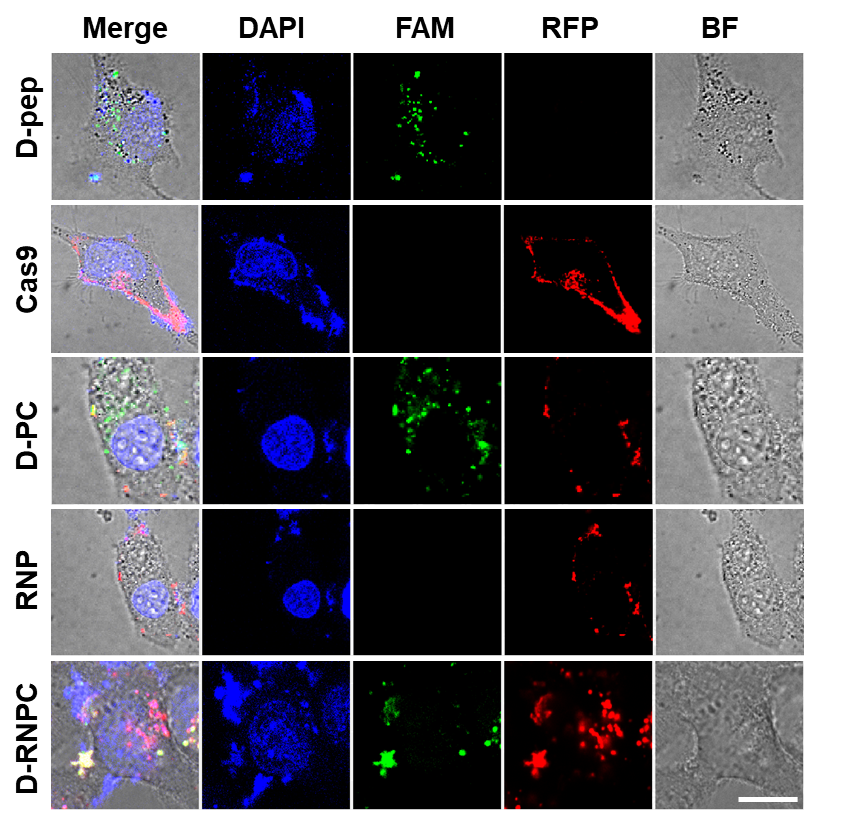

**Figure S13.** Cellular uptake of D-RNPC. Confocal images of HeLa cell incubated with 200 nM D-RNPC, RNP, D-PC, Cas9 and 200 µM D-pep. Scale bar, 25 µm.

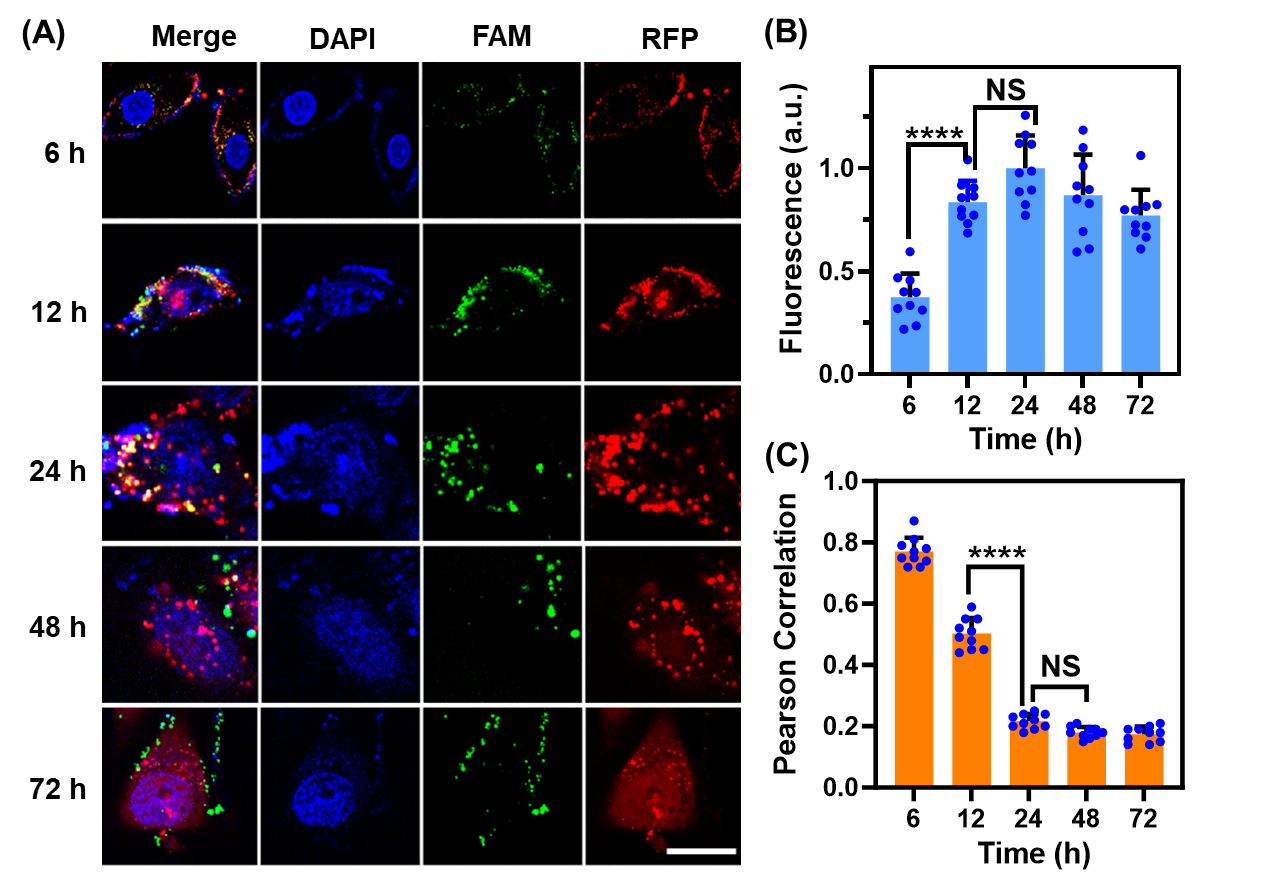

**Figure S14.** D-RNPC internalization times optimization. (**A**) Confocal images of HeLa cells incubated with 200 nM D-RNPC for different times. (**B**) The fluorescence intensity of FAM in confocal images. (**C**) The pearson correlation analysis was performed on the FAM and RFP in confocal images. Scale bar, 25 µm. Shown are the mean and the standard error of the mean (SEM) values from ten individual cells. Two-tailed Student's t-test: ****P < 0.0001.

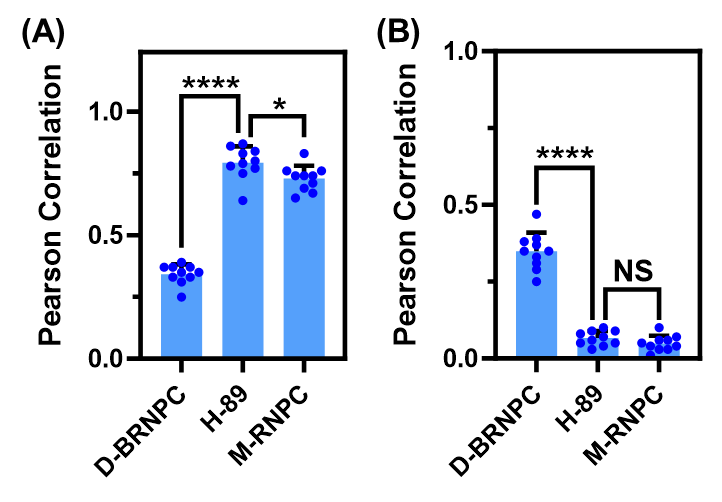

**Figure S15.** Pearson correlation analysis of Figure. 3a (**A**) The pearson correlation analysis was performed on the peptide (green) and Cas9 (red) in confocal images. Shown are the mean and the standard error of the mean (SEM) values from ten individual cells. Two-tailed Student's t-test: *P < 0.1. ****P < 0.0001. (**B**) The pearson correlation analysis was performed on the cell nucleus (blue) and Cas9 (red) in confocal images. Shown are mean ± SEM from ten individual cells. Two-tailed Student's t-test: ****P<0.0001.

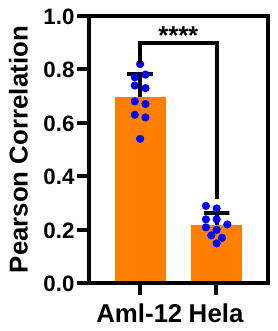

**Figure S16.** Pearson correlation analysis of FAM and RFP in Figure. 3b Shown are the mean and the standard error of the mean (SEM) values from ten individual cells. Two-tailed Student's t-test. ****P < 0.0001.

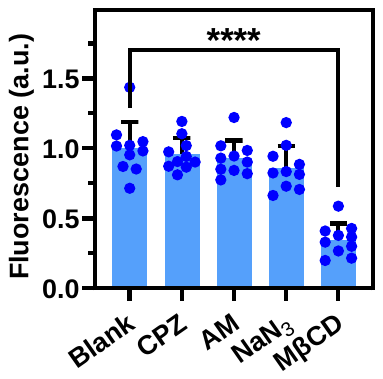

**Figure S17.** Analysis the fluorescence intensity of RFP from Figure. 3d Shown are the mean and the standard error of the mean (SEM) values from ten individual cells. Two-tailed Student's t-test: ****P < 0.0001.

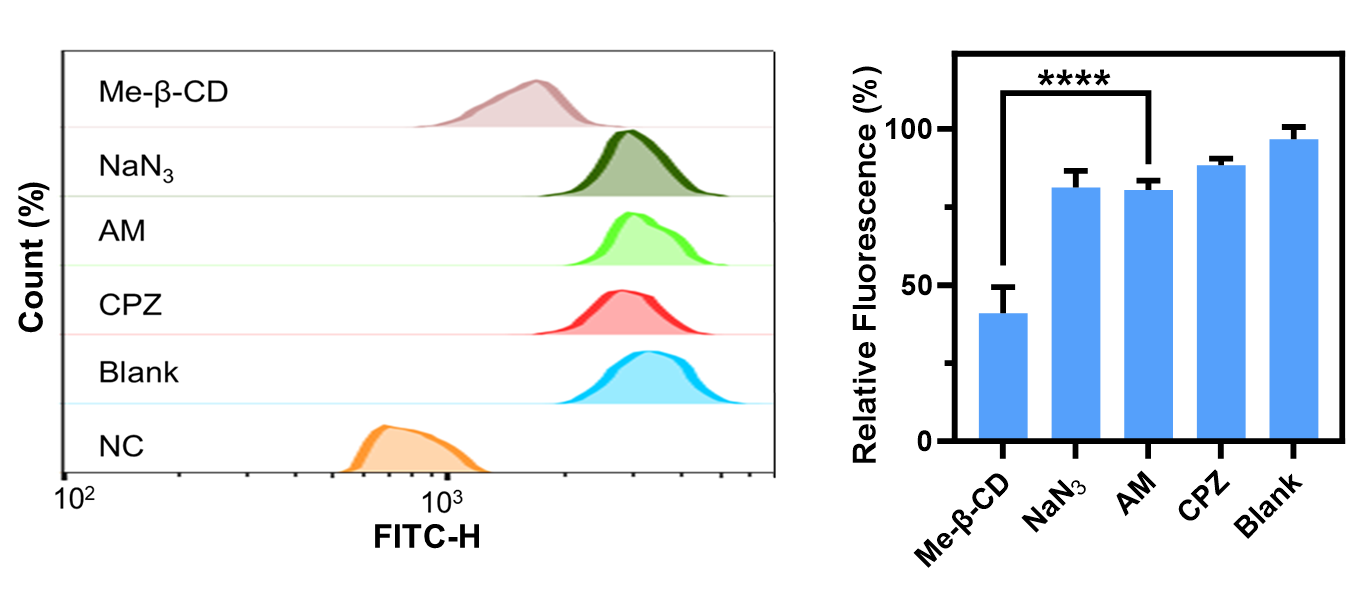

**Figure S18.** Cellular internalization pathways of D-RNPC. Flow cytometry and fluorescence intensity analysis of HeLa cells pretreated without (Blank) and with Chlorpromazine (CPZ), Amiloride (AM), Methyl-β-cyclodextrin (Me-β-CD), Sodium azide (NaN3), then incubated with and without (NC) D-RNPC. Shown are the mean and the standard error of the mean (SEM) values from three experiments. Two-tailed Student's t-test: ****P < 0.0001.

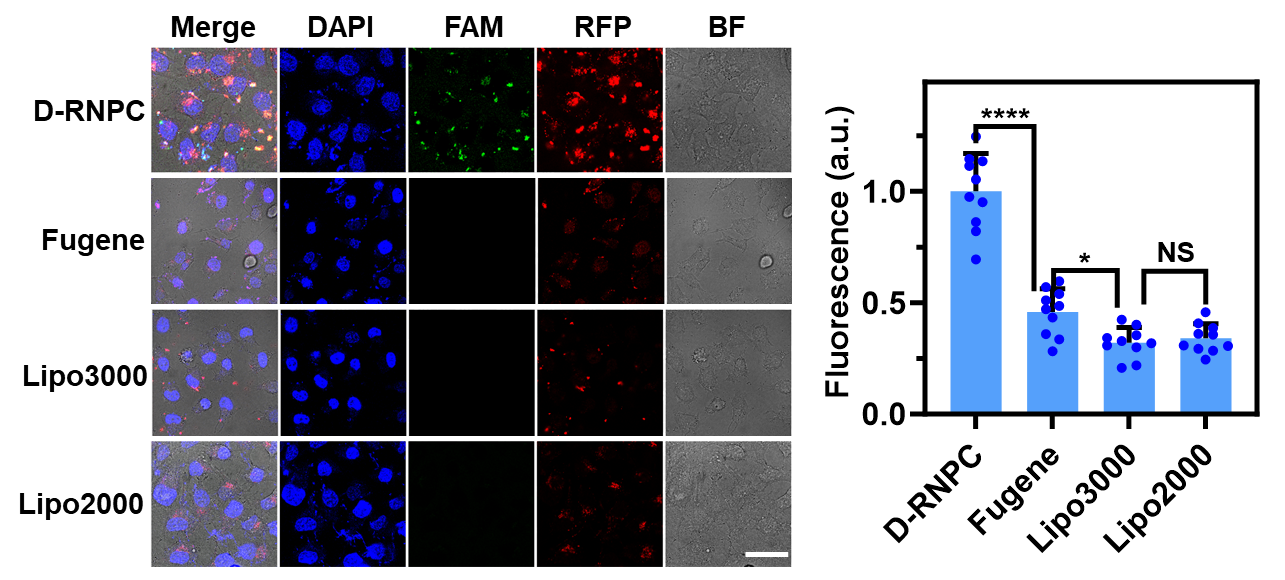

**Figure S19.** Comparison of transfection efficiency. Confocal images and RFP fluorescence intensity analysis of HeLa cells incubated with 200 nM D-RNPC. As control, the cells were transfected 200 nM RNP with Lipo2000, Lipo3000 and Fugene. Scale bar, 60 µm. Shown are the mean and the standard error of the mean (SEM) values from ten individual cells. Two-tailed Student's t-test: ****P<0.0001, *P<0.1.

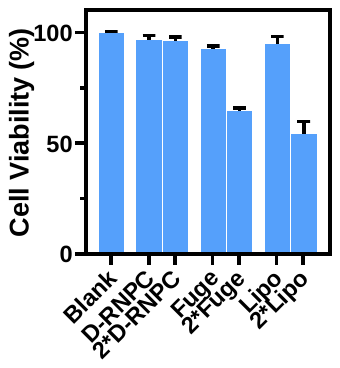

**Figure S20.** Comparison of cytotoxicity. Cytotoxicity induced without (Blank) and with 200 nM D-RNPC, 400 nM D-RNPC (2*D-RNPC), 1 uL FugeneHD transfection of 200 nM RNP (Fuge), 2 uL FugeneHD transfection of 400 nM RNP (2*Fuge), 1 uL Lipo 2000 transfection of 200 nM RNP (Lipo), and 2 uL Lipo 2000 transfection of 400 nM RNP (2*Lipo) in 12 h incubation. Shown are the mean and the standard error of the mean (SEM) values from three individual experiments.

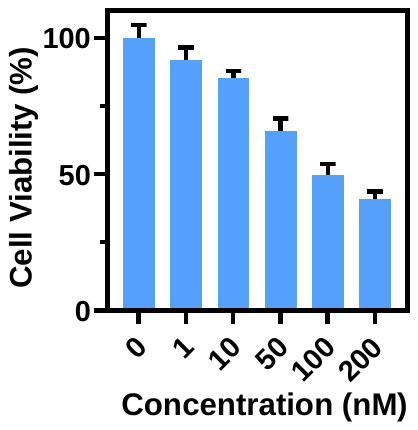

**Figure S21.** Cytotoxicity of D-RNPC in HeLa cells. Cell viability of HeLa cells treated with different concentrations of D-RNPC (0 nM, 1 nM, 10 nM, 50 nM, 100 nM, 200 nM) for 72 h. Shown are the mean and the standard error of the mean (SEM) values from three experiments.

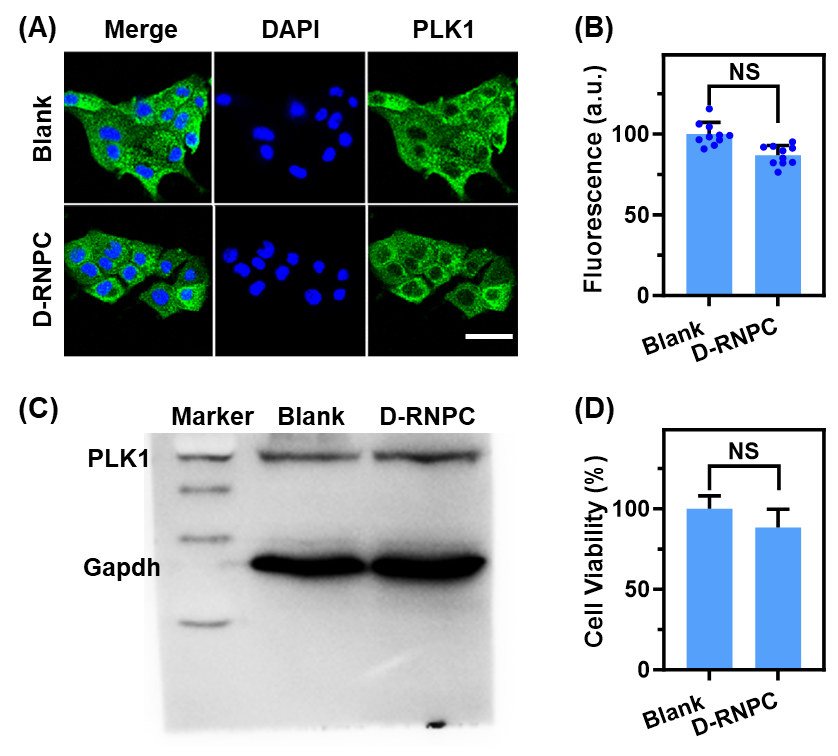

**Figure S22.** *In vitro* gene editing and therapy of D-RNPC in AML-12 cells. Immunofluorescence analysis (**A**) and Western blotting analysis (**C**) of PLK1 protein in AML-12 cells treated without (Blank) and with 200 nM D-RNPC for 72 h. Scale bar, 50 µm. (**B**) Statistical analysis of PLK1 fluorescence intensity in Figure S22A. Shown are mean ± SEM from ten individual cells. (**D**) Cell viability of AML-12 cells treated without (Blank) and with 200 nM D-RNPC for 72 h. Shown are mean ± SEM from three experiments.

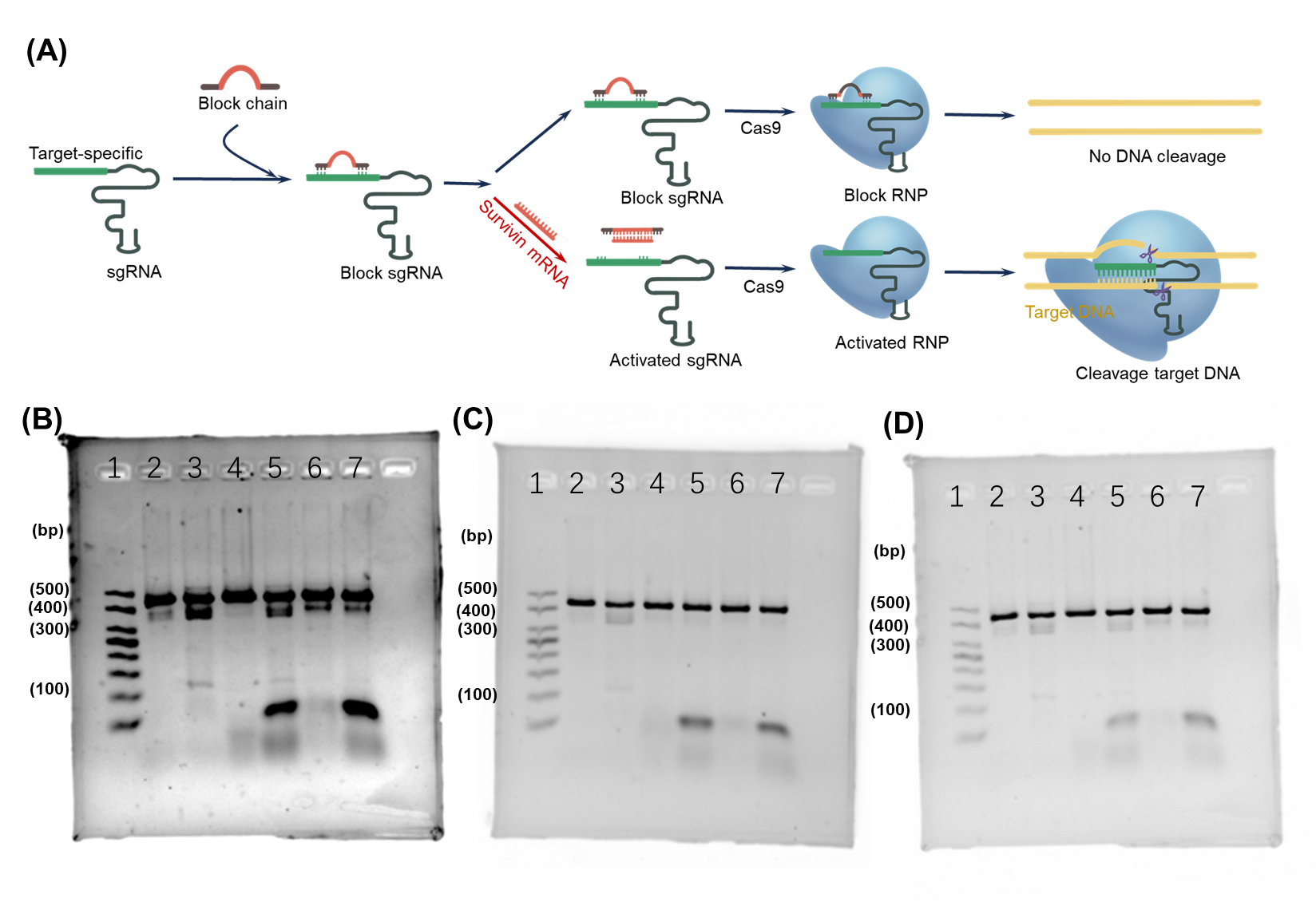

**Figure S23.** Design and optimization of F-D-BRNPC system. (**A**) Schematic diagram of sgRNA blockade and activation (**B**) 1. DNA ladder marker; 2. DNA; 3. mixture of DNA and RNP; 4. mixture of DNA and block 1 RNP; 5. mixture of DNA, block 1 RNP and target mRNA; 6. mixture of DNA and block 2 RNP; 7. mixture of DNA, block 2 RNP and target mRNA. (**C**) 1. DNA ladder marker; 2. DNA; 3. mixture of DNA and RNP; 4. mixture of DNA and block 3 RNP; 5. mixture of DNA, block 3 RNP and target mRNA; 6. mixture of DNA and block 4 RNP; 7. mixture of DNA, block 4 RNP and target mRNA. (**D**) 1. DNA ladder marker; 2. DNA; 3. mixture of DNA and RNP; 4. mixture of DNA and block 5 RNP; 5. mixture of DNA, block 5 RNP and target mRNA; 6. mixture of DNA and block 6 RNP; 7. mixture of DNA, block 6 RNP and target mRNA.

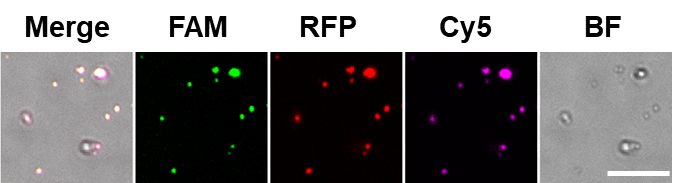

**Figure S24.** Confocal images of F-D-BRNPC. FAM, RFP and Cyanine 5 (cy5) were labeled peptides, cas9, and block chain1, respectively. Scale bar, 5 µm.

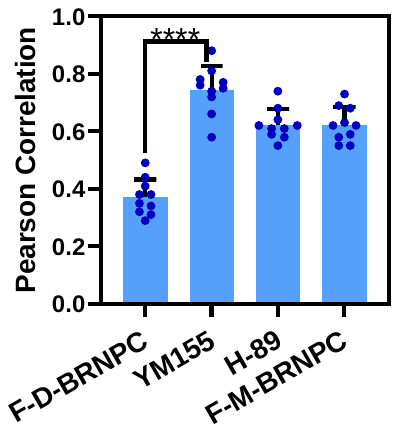

**Figure S25.** Pearson correlation coefficient analysis the red and purple fluorescence signals in Figure. 4a Shown are the mean and the standard error of the mean (SEM) values from ten individual cells. Two-tailed Student's t-test. ****P < 0.0001.

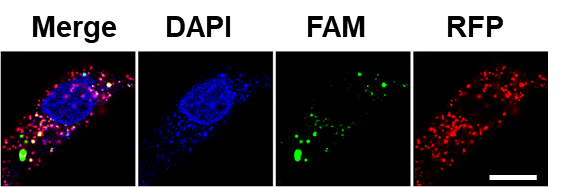

**Figure S26.** Cellular internalization of F-D-BRNPC. Confocal fluorescence microscopic images of HeLa cell incubated with 200 nM F-D-BRNPC. Scale bar, 25 µm.

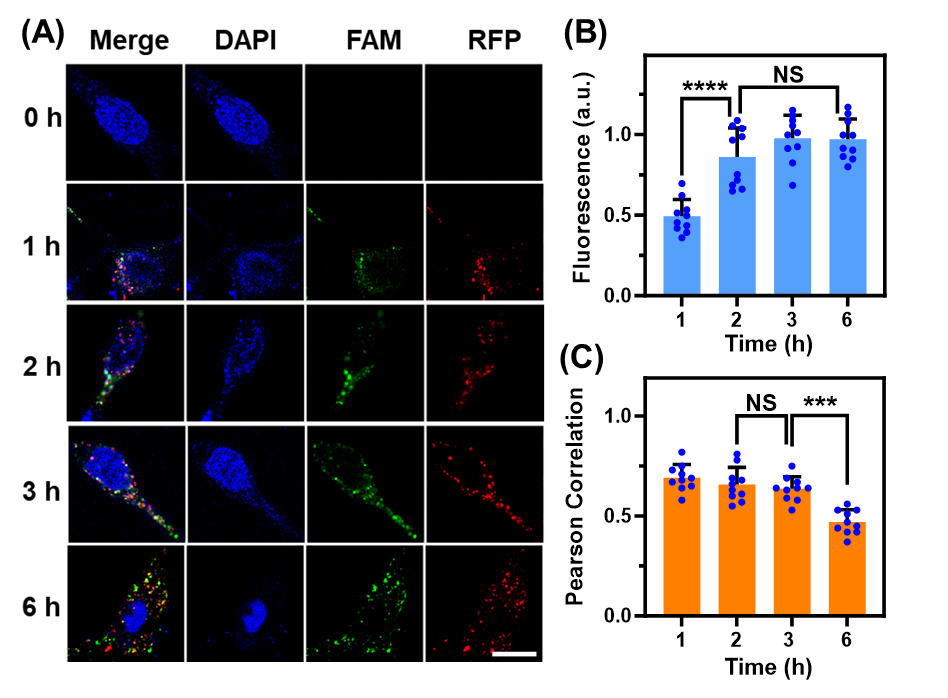

**Figure S27.** Times optimization of FA-D-BRNPC in HeLa cells. (**A**) Confocal images (**B**) FAM fluorescence intensity statistics (**C**) RFP and FAM Pearson coefficient analysis of HeLa cells. Scale bar, 25 µm. Shown are the mean and the standard error of the mean (SEM) values from ten individual cells. Two-tailed Student's t-test. ***P < 0.001, ****P < 0.0001.

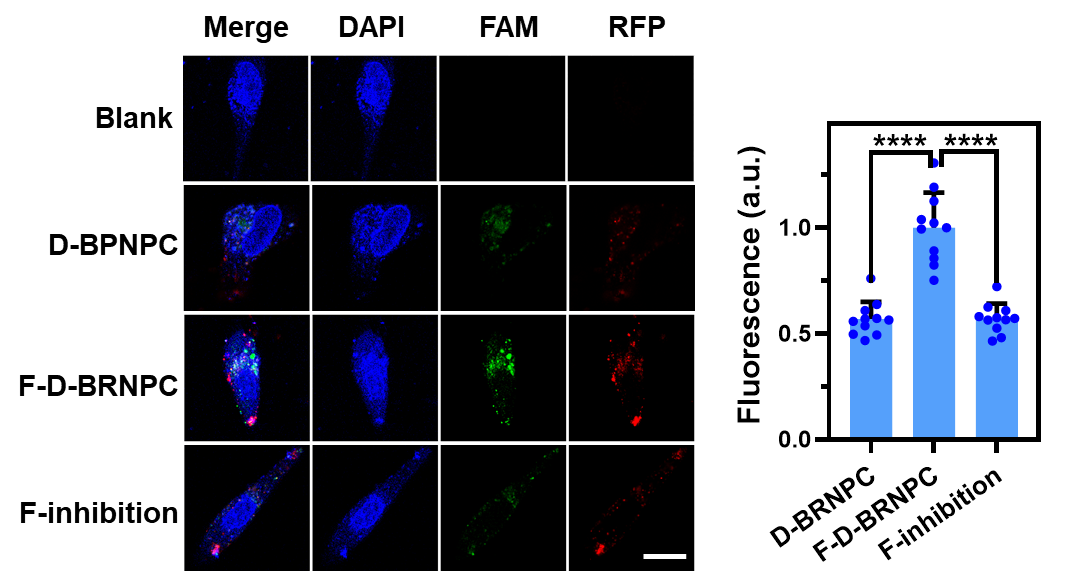

**Figure S28.** Cell targeting of F-D-BRNPC. Confocal images and FAM fluorescence intensity statistics of HeLa cells incubated without (Blank) and with 200 nM D-BRNPC, 200 nM F-D-BRNPC in absence and presence of 1 µM folic acid (F-inhibition) for 2 h. Scale bar, 25 µm. Shown are the mean and the standard error of the mean (SEM) values from ten individual cells. Two-tailed Student's t-test. ****P < 0.0001.

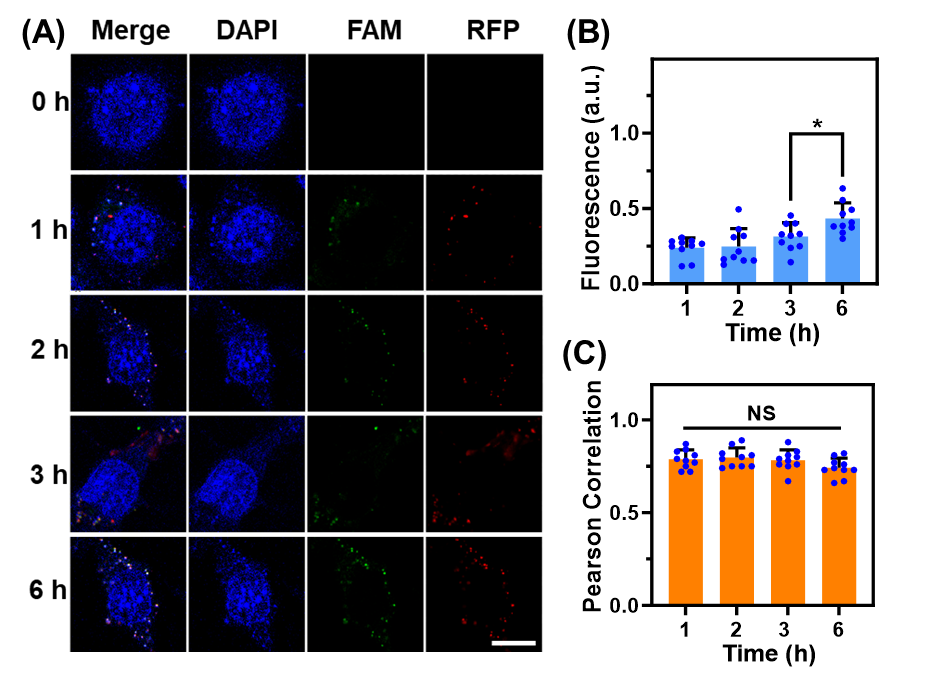

**Figure S29.** Times optimization of FA-D-BRNPC in AML-12 cells. (**A**) Confocal images (**B**) FAM fluorescence intensity statistics (**C**) RFP and FAM Pearson coefficient analysis of AML-12 cells. Scale bar, 25 µm. Shown are the mean and the standard error of the mean (SEM) values from ten individual cells. Two-tailed Student's t-test. *P < 0.1.

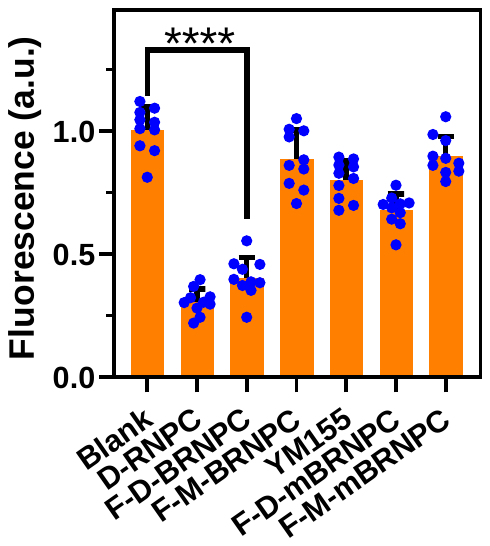

**Figure S30.** Analysis of PLK1 fluorescence intensity (Green) in Figure. 5c Shown are the mean and the standard error of the mean (SEM) values from ten individual cells. Two-tailed Student's t-test. ****P<0.0001.

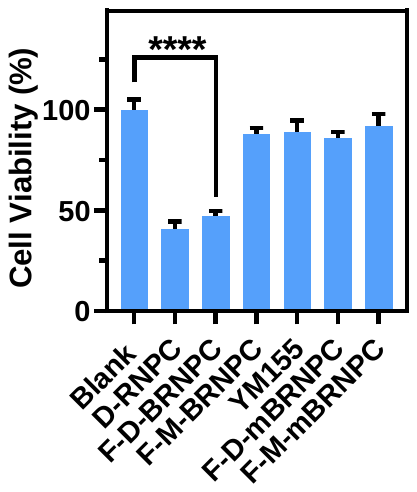

**Figure S31.** Cell viability of HeLa cells induced with F-D-BRNPC. Cell viability of HeLa cells treated without (Blank) and with 200 nM D-RNPC, 200 nM F-D-BRNPC, 200 nM F-M-BRNPC, 200 nM F-D-BRNPC and 10 nM YM155 (YM155) mixture, 200 nM F-D-mBRNPC, or 200 nM F-M-mBRNPC for 72 h. Shown are mean ± SEM from three experiments. ****P<0.0001.

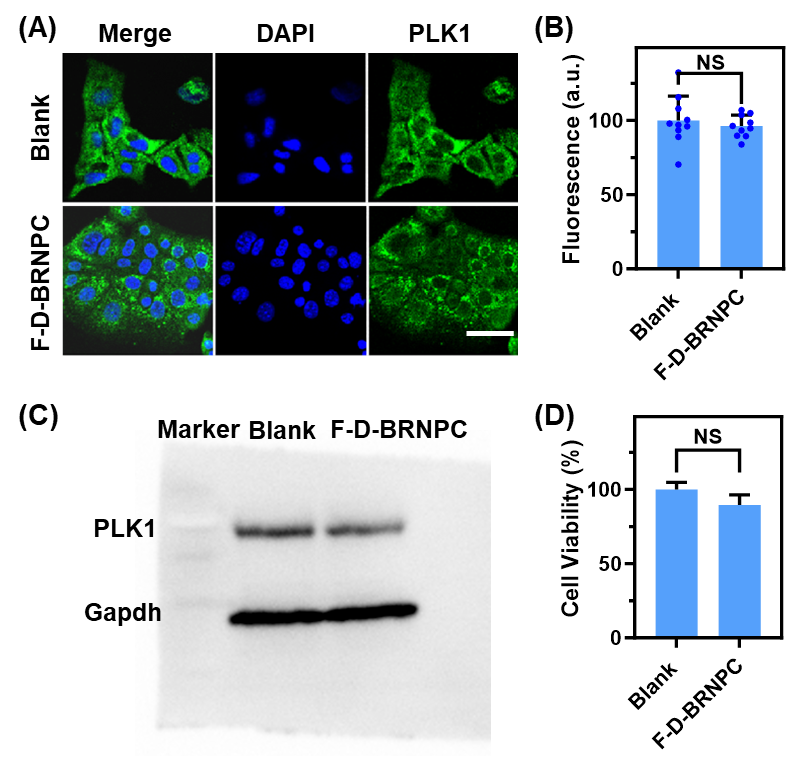

**Figure S32.** *In vitro* gene editing and therapy of D-BRNPC in AML-12 cells. Immunofluorescence analysis (**A**) and Western blotting analysis (**C**) of PLK1 protein in AML-12 cells treated without (Blank) and with 200 nM F-D-BRNPC for 72 h. Scale bar, 50 µm. (**B**) Statistical analysis of PLK1 fluorescence intensity in Figure S22A. Shown are mean ± SEM from ten individual cells. (**D**) Cell viability of AML-12 cells treated without (Blank) and with 200 nM F-D-BRNPC for 72 h. Shown are mean ± SEM from three experiments.

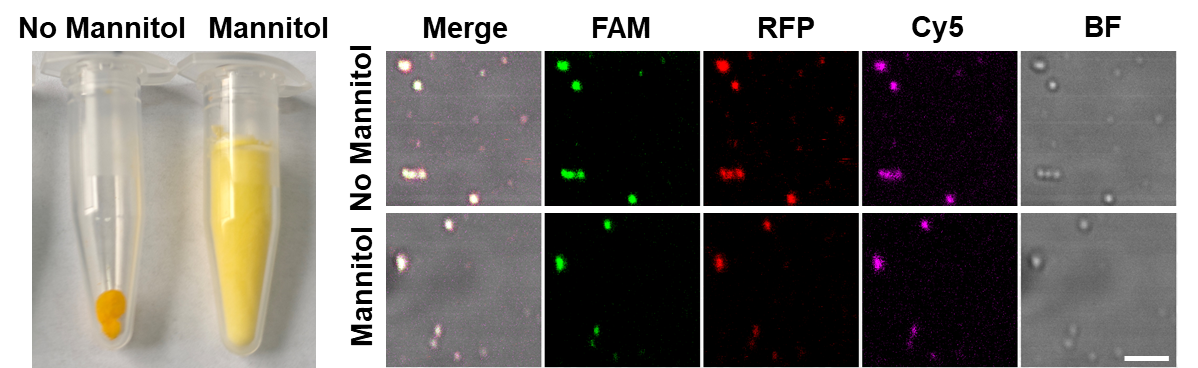

**Figure S33.** Morphology and confocal images of lyophilized F-D-BRNPC without and with 1.5% mannitol. Scale bar, 5 µm.

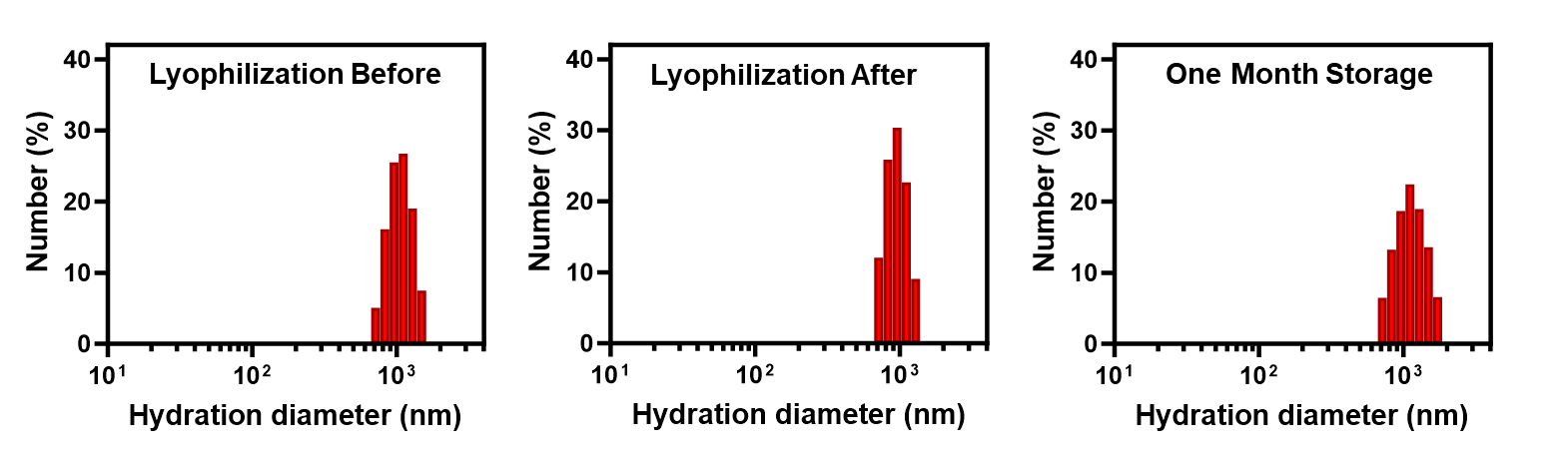

**Figure S34.** DLS analysis of hydrodynamic diameter distribution of F-D-BRNPC before and after lyophilization and lyophilized F-D-BRNPC with one month storage.

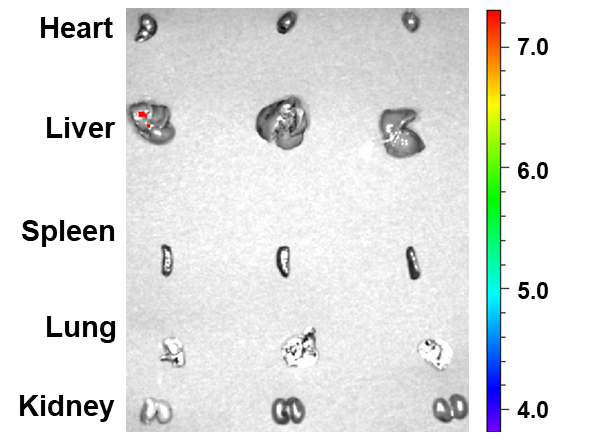

**Figure S35.** *Ex vivo* images of mouse major organs that received PBS buffer after 2  h inhalation administration.

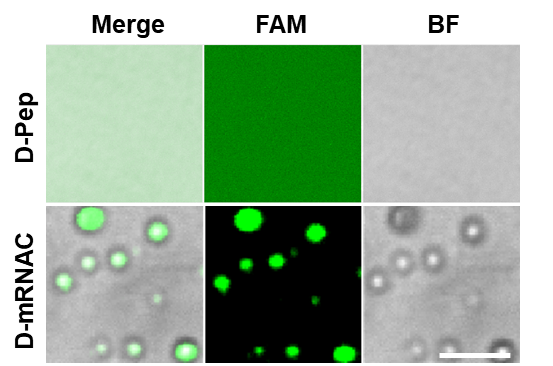

**Figure S36.** Confocal images of D-mRNAC and D-Pep. Scale bar, 5 µm.

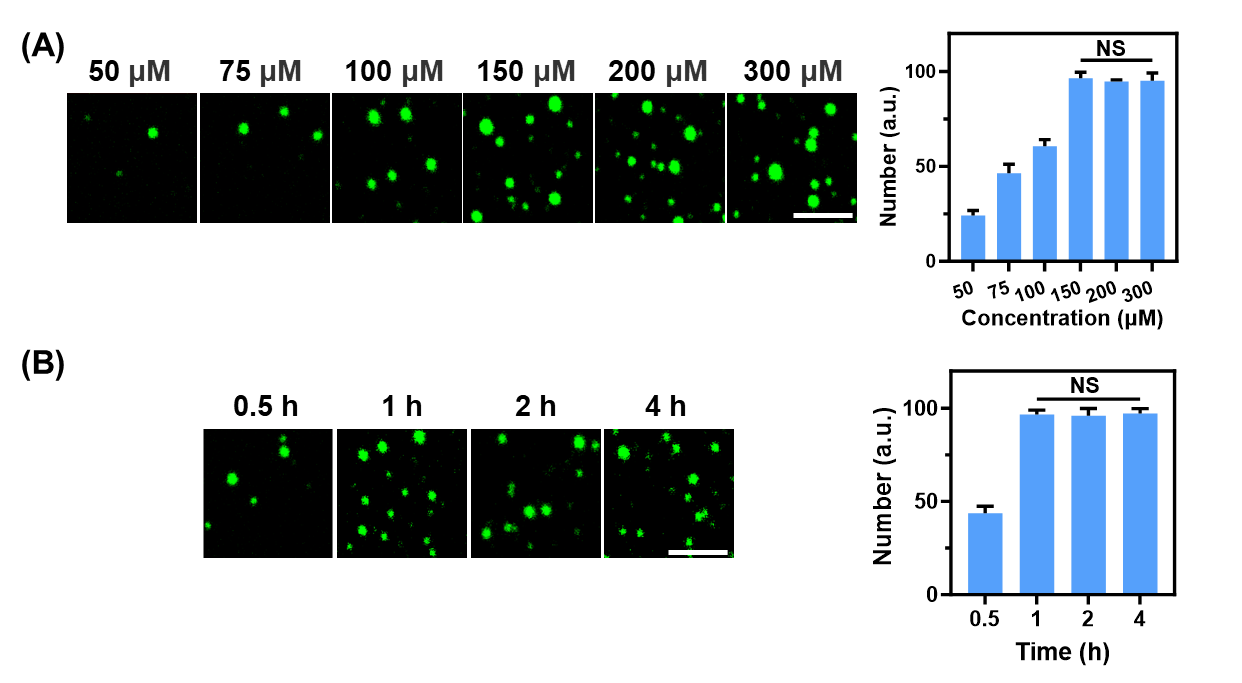

**Figure S37.** Confocal images and number statistics of D-mRNAC. Optimization of peptide concentrations (**A**) and incubation times (**B**) for D-mRNAC preparation. Shown are the mean and the standard error of the mean (SEM) values from at least three representative images.

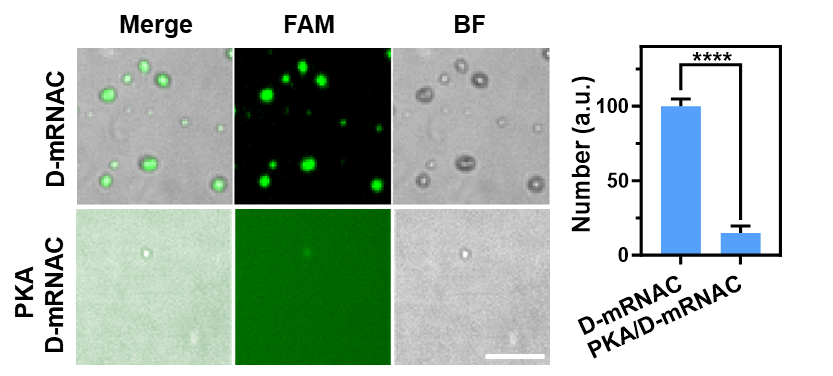

**Figure S38.** PKA triggered disassembly of D-mRNAC. Confocal images and number statistics of 200 nM D-mRNAC before (D-mRNAC) and after (PKA-D-mRNAC) addition of 1400 nM PKA for 1 h incubation. Scale bar, 5 µm. Shown are the mean and the standard error of the mean (SEM) values from at least three representative images. Two-tailed Student's t-test. ****P<0.0001.

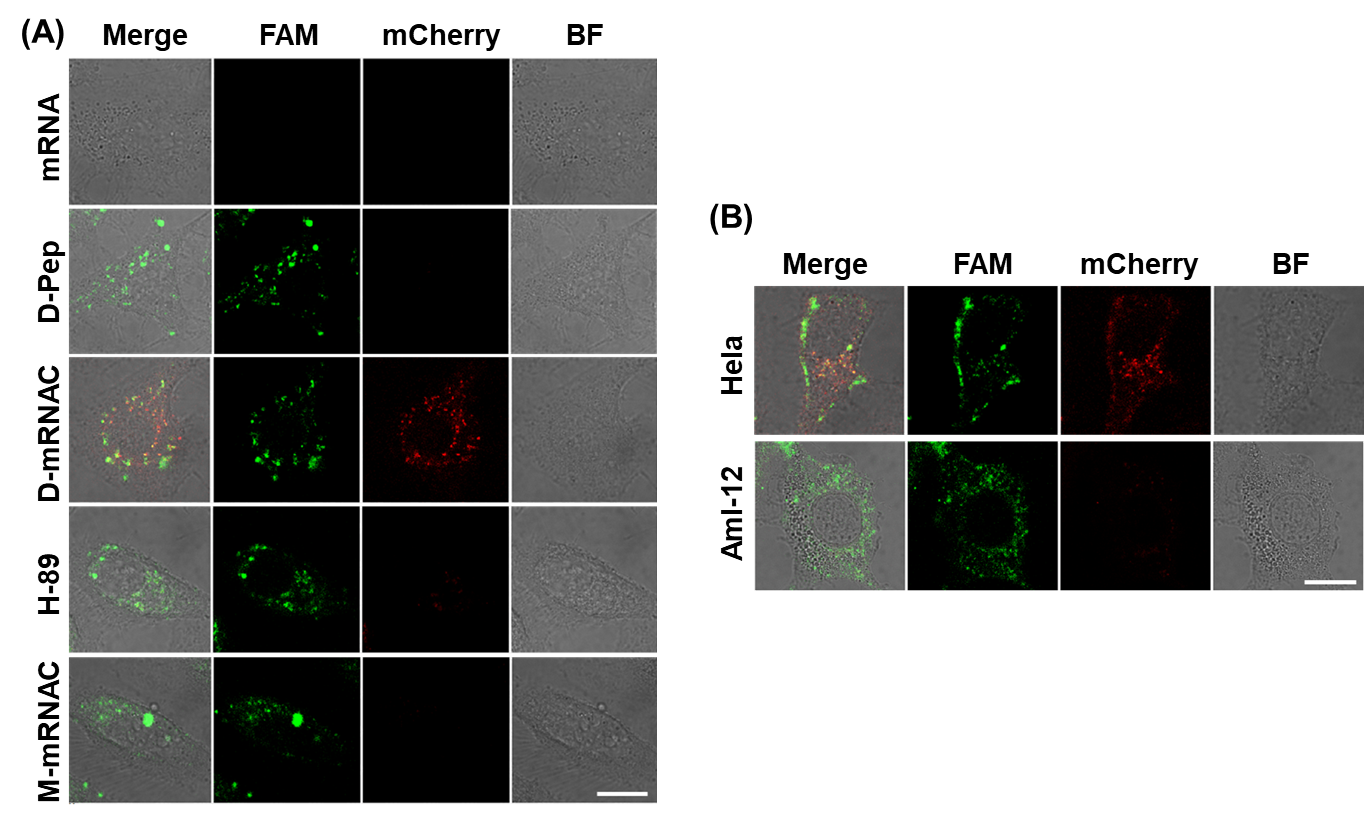

**Figure S39.** Performance of D-mRNAC in cells. (**A**) Confocal images of HeLa cells incubated with 150 µM D-Pep (D-Pep), 200 nM D-mRNAC (D-mRNAC), 200 nM D-mRNAC with 20 μM H-89 (H-89), or 200 nM M-mRNAC (M-mRNAC). Scale bar, 25 µm. (**B**) Confocal images of HeLa cells and AML-12 cells incubated with 200 nM D-mRNAC. Scale bar, 25 µm.

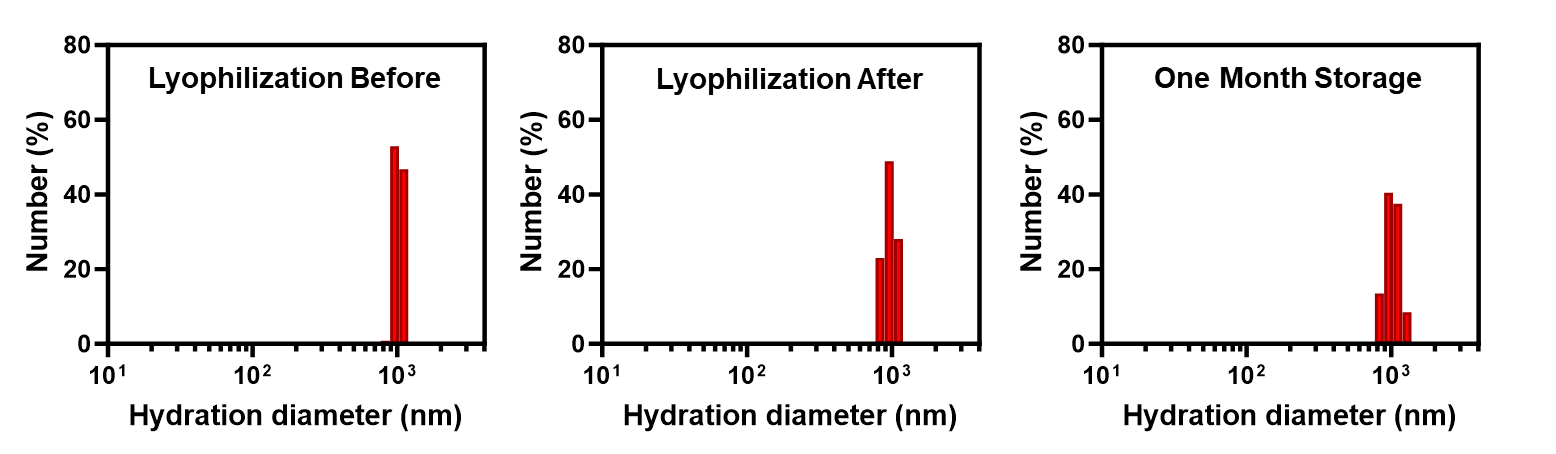

**Figure S40.** DLS analysis of hydrodynamic diameter distribution of D-mRNAC before and after lyophilization and lyophilized D-mRNAC with one month storage.

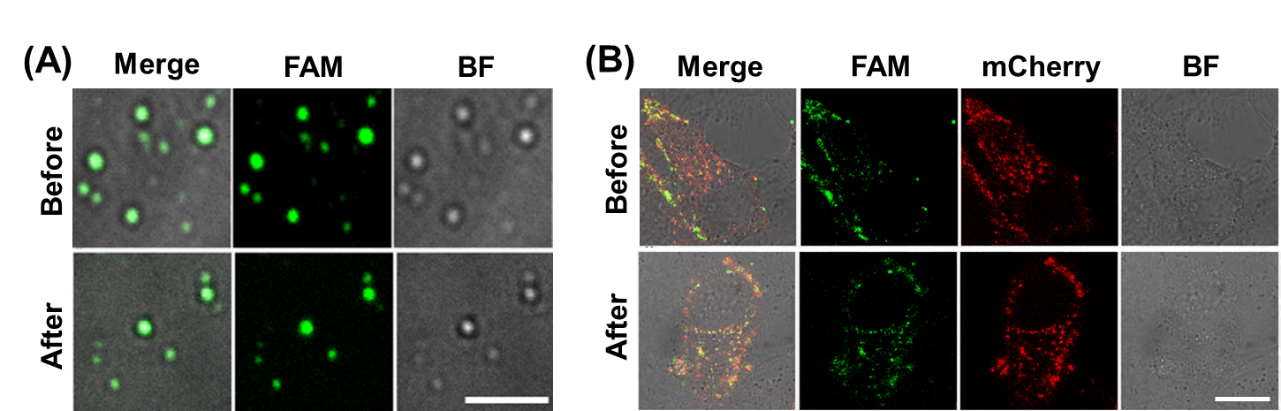

**Figure S41.** Comparison of D-mRNAC before and after lyophilization. (**A**) Confocal images of D-mRNAC before and after lyophilization. Scale bar, 5 µm. (**B**) Confocal images of HeLa cells incubated with 200 nM D-mRNAC or lyophilized D-mRNAC. Scale bar, 25 µm.

**Table S1.** Oligonucleotide sequences used in this study.

| **Names** | **Sequences from 5’ to 3’** |
| --- | --- |
| Dual peptide | RRASLRRASL |
| Dual peptide | 5-FAM-RRASLRRASL |
| Triple peptide | 5-FAM-RRASLRRASLRRASL |
| Misorder peptide | 5-FAM-RSRALRSRAL |
| sgRNA (PLK1) | GGCCGACCCTGGGAAAGCCG |
| Block chain 1 | TTTCCCGAATGTAGAGATGCGGTGGGGGTC |
| Block chain 2 | CTTTCCCGAATGTAGAGATGCGGTGGGGGTCG |
| Block chain 3 | GCTTTCCCGAATGTAGAGATGCGGTGGGTCGG |
| Block chain 4 | GCTTTCCCGAATGTAGAGATGCGGTGGGTCGGC |
| Block chain 5 | AAACCGGCGAATGTAGAGATGCGGTGGTCGGCC |
| Block chain 6 | AACCGGCGAATGTAGAGATGCGGTGGTCGGC |
| Misorder Block chain 1 | TTTCCCGATGTGGAAGTCGAGTGAGGGGTC |
| Folic acid Block chain 1 | Folic acid-TTTCCCGAATGTAGAGATGCGGTGGGGGTC-CY5 |
| *Survivin* mRNA | CCACCGCATCTCTACATTC |
| F-T7E1 | GGAGGCTCTGCTCGGATCGA |
| R-T7E1 | CAACACCACGAACACGAAGT |

The sequences of Survivin mRNA and the complementary regions are shown in red
